# mTORC2 stabilizes HIF-1β to coordinate metabolic adaptation in lung cancer

**DOI:** 10.64898/2026.09.02.748876

**Authors:** Morgane Paque, Ning An, Marine Leclercq, Anisa Aliju, Elodie Renaude, Domien Vanneste, Denis Heusdens, Jair Marques Junior, Chinmayi Pednekar, Tillmann Bedau, Juan Fernández-García, Fabienne Perin, Raphaël Vanleyssem, Christian Seca, Kim Eiden, Patrick Roncarati, Engy Shokry, Rachel L. Baird, Leah Officer-Jones, Silvia Martinelli, Ian Powley, David Sumpton, Thomas Marichal, Didier Cataldo, Emmanuel di Valentin, Steven Goosens, Geert Berx, John Le Quesne, Michael Herfs, Johannes Meiser, Reinhard Büttner, Mélanie Planque, Johannes V. Swinnen, Sarah-Maria Fendt, Alex von Kriegsheim, Pierre Close, Arnaud Blomme

## Abstract

Despite extensive genetic heterogeneity, lung tumors frequently converge on shared signaling dependencies that remain therapeutically underexploited. Here, we identify mTORC2 signaling as a convergent dependency across genetically distinct lung cancer subtypes and uncover HIF-1β as a selective metabolic effector downstream of mTORC2 that promotes lung tumor progression. Elevated mTORC2 signaling in lung adenocarcinoma was associated with poor overall survival, metastatic dissemination and metabolic rewiring. Using complementary genetically engineered mouse models of *Rictor* deletion or overexpression in *Kras*-driven lung tumors, we show that mTORC2 activity is dispensable for normal lung homeostasis but required for tumor progression and metabolic adaptation *in vivo*. Mechanistically, mTORC2 stabilized HIF-1β by preventing its ubiquitin-independent proteasomal degradation through a non-canonical PKCα-CK2 signaling axis, independently of AKT. Integrated multi-omics analyses identified extensive metabolic rewiring downstream of the mTORC2-HIF-1β axis, with sphingolipid metabolism emerging as a prominent and therapeutically exploitable vulnerability. Accordingly, pharmacological targeting of sphingolipid metabolism markedly impaired the growth of mTORC2-driven lung tumors *in vivo*. Together, our findings establish a non-canonical mTORC2-HIF-1β signaling axis that couples oncogenic signaling to metabolic adaptation and defines therapeutically actionable metabolic vulnerabilities in lung cancer.

## Introduction

Lung cancer remains the leading cause of cancer-related mortality worldwide, with non-small cell lung cancer (NSCLC) accounting for approximately 85% of cases (1). Despite major therapeutic advances, durable clinical benefit remains limited by the extensive genetic, epigenetic, metabolic and phenotypic heterogeneity of NSCLC (2–4). The identification of actionable driver mutations has enabled the development of targeted therapies that have significantly improved patient outcome, including in *EGFR*-, *ALK*-and *KRAS^G12C^*^-^mutant tumors (5). However, resistance frequently emerges through secondary genetic alterations and adaptive signaling rewiring (6,7). Notably, despite their molecular diversity, lung tumors often converge on shared signaling dependencies that sustain tumor progression (8,9). These observations suggest that targeting convergent adaptive dependencies, rather than individual oncogenic drivers, may represent a more durable therapeutic strategy in lung cancer.

Aberrant activation of the PI3K-AKT-mTOR pathway is a common feature of NSCLC and has therefore prompted extensive efforts to therapeutically target mTOR signaling (10). The serine/threonine kinase mTOR assembles into two structurally and functionally distinct complexes: mTORC1, defined by RAPTOR, and mTORC2, characterized by RICTOR (11). While therapeutic strategies have historically focused on mTORC1, increasing evidence suggests that mTORC2 may represent a particularly important adaptive signaling node in lung cancer. Indeed, recurrent *RICTOR* amplification and elevated RICTOR expression are frequently observed in NSCLC and are associated with poor prognosis (12,13). In contrast to mTORC1, which primarily controls anabolic growth programs, mTORC2 functions as a context-dependent survival and metabolic adaptation pathway that becomes critical under oncogenic and therapeutic stress (14). Moreover, mTORC2 signaling can persist following mTORC1 inhibition, thereby sustaining adaptive tumor signaling through AKT activation. These observations may explain the limited efficacy of rapamycin analogues and other mTOR-targeted therapies and raise the possibility that mTORC2-directed strategies could provide greater therapeutic benefit in lung cancer. However, the development of selective pharmacological inhibitors of mTORC2 remains challenging (15), highlighting the need to identify therapeutically actionable downstream vulnerabilities associated with mTORC2 activation.

Beyond its canonical role in growth and survival signaling, mTORC2 has emerged as a central regulator of cancer metabolism. Previous studies have linked mTORC2 to glucose utilization, glycolytic metabolism and cellular stress adaptation, suggesting that it coordinates metabolic programs required for tumor progression under adverse microenvironmental conditions (16–18). HIF-dependent transcriptional programs are central mediators of metabolic adaptation, yet whether and how mTORC2 engages such programs remains poorly understood (19). In parallel, increasing evidence implicates mTORC2 in the control of lipid metabolism (20), including de novo lipogenesis and sphingolipid metabolism, processes that contribute to tumor progression in multiple cancer types (21,22). In addition to their structural role in membrane organization, sphingolipids function as bioactive signaling molecules regulating proliferation, stress adaptation and immune microenvironment remodeling (23). Consistent with these observations, enzymes involved in sphingolipid homeostasis are frequently dysregulated in NSCLC and have been associated with therapeutic resistance and poor clinical outcome (24–27). Together, these observations suggest that identifying the metabolic programs engaged downstream of mTORC2 may reveal novel therapeutic vulnerabilities in lung cancer.

Here, we identify mTORC2 signaling as a convergent adaptive dependency in lung cancer and demonstrate that RICTOR-dependent mTORC2 activity promotes lung cancer progression through HIF-1β-dependent metabolic rewiring. Using complementary gain-and loss-of-function genetically engineered mouse models (GEMMs) together with integrated proteomic, transcriptomic and lipidomic analyses and human patient datasets, we uncover a non-canonical mTORC2-HIF-1β signaling axis that regulates lipid metabolism in lung tumors. Mechanistically, mTORC2 stabilizes HIF-1β through a PKCα-CK2-dependent pathway, independently of AKT, thereby promoting lipid metabolic programs associated with tumor progression. Importantly, pharmacological targeting of sphingolipid metabolism markedly impaired the growth of mTORC2-driven lung tumors, identifying targetable metabolic vulnerabilities downstream of mTORC2 signaling in lung cancer.

## Results

### mTORC2 activation is a recurrent feature of aggressive lung tumors

Lung tumors are characterized by extensive genetic heterogeneity that contributes to therapeutic resistance and disease progression. Identifying signaling pathways that are recurrently engaged across genetically distinct tumors may therefore uncover targetable adaptive dependencies. To identify signaling networks associated with oncogene-driven lung tumorigenesis, we performed proteomics on *Kras^G12D^*^-^driven lung tumors and control lung tissues (isolated from *Ccsp-Cre;Kras^G12D/+^* and *Ccsp-Cre;Kras^+/+^* mice, further referred to as *Ccsp-Kras^G12D/+^* and *Ccsp* mice respectively) (Fig. 1A, Fig. S1A). Gene Set Enrichment Analysis (GSEA) revealed significant enrichment of metabolic and stress adaptation programs in tumor-bearing lungs, including oxidative phosphorylation, glycolysis, lipid metabolic pathways and mTOR signaling (Fig. 1B). Strikingly, *Kras*-driven lung tumors selectively accumulated RICTOR protein, whereas RAPTOR expression was only modestly increased (Fig. 1C-D). In parallel, *Kras-*mutant tumors exhibited increased mTOR expression and phosphorylation on Ser2448 (Fig. 1D), together with enhanced phosphorylation of RAPTOR on Ser792, indicative of AMPKα activation, and reduced phosphorylation of RICTOR on Thr1135 (Fig. S1B). Collectively, these findings support preferential activation of mTORC2 signaling in *Kras*-driven lung tumors. In human lung adenocarcinoma (LUAD), genomic amplification of *RICTOR* (*RICTOR^amp^*^)^ is frequently observed and is often considered as the only relevant genomic alteration within those tumors (13). Analysis of The Cancer Genome Atlas (TCGA) LUAD cohort confirmed that *RICTOR^amp^* occurs in >8% of lung adenocarcinoma (43/507, Fig. 1E) and correlates with increased RICTOR protein abundance, without affecting RAPTOR expression (Fig. 1F-G). Notably, despite representing three distinct molecular subsets of lung cancer, *RICTOR^amp^*, *KRAS^mut^* and *EGFR^mut^* tumors all exhibited elevated RICTOR protein expression across multiple datasets. In contrast, RAPTOR protein expression remained comparatively unchanged in these molecular subsets (Fig. 1F-K, Fig. S1C-F). These findings suggest that distinct oncogenic alterations may functionally converge toward elevated mTORC2 signaling in lung cancer. To further assess mTORC2 activity in human lung tumors, we performed multiplexing immunofluorescence and immunohistochemistry (IHC) on LUAD biopsies. In two independent patient cohorts, RICTOR expression positively correlated with phosphorylation of AKT on Ser473, consistent with active mTORC2 signaling in lung tumors (r = 0.7268 and r = 0.5008; Fig. 1L-O, Fig. S1G). High levels of RICTOR (RIC^high^ tumors) were further associated with increased lymph node infiltration and distant metastasis dissemination (36% of lymph node involvement and 66.7% of distant metastasis in RIC^high^ tumors versus 20.4% and 27.8% in RIC^low^ tumors respectively, Fig. 1P-Q). Moreover, elevated intratumoral RICTOR expression at diagnosis was associated with an increased risk of developing metastasis (Fig. 1R). Finally, high RICTOR levels significantly correlated with shortened patient survival (Fig. 1S). Together, these results identify mTORC2 activation as a recurrent feature across molecularly distinct lung tumors and support a clinically relevant role for RICTOR-dependent signaling in lung cancer progression.

**Figure 1:**
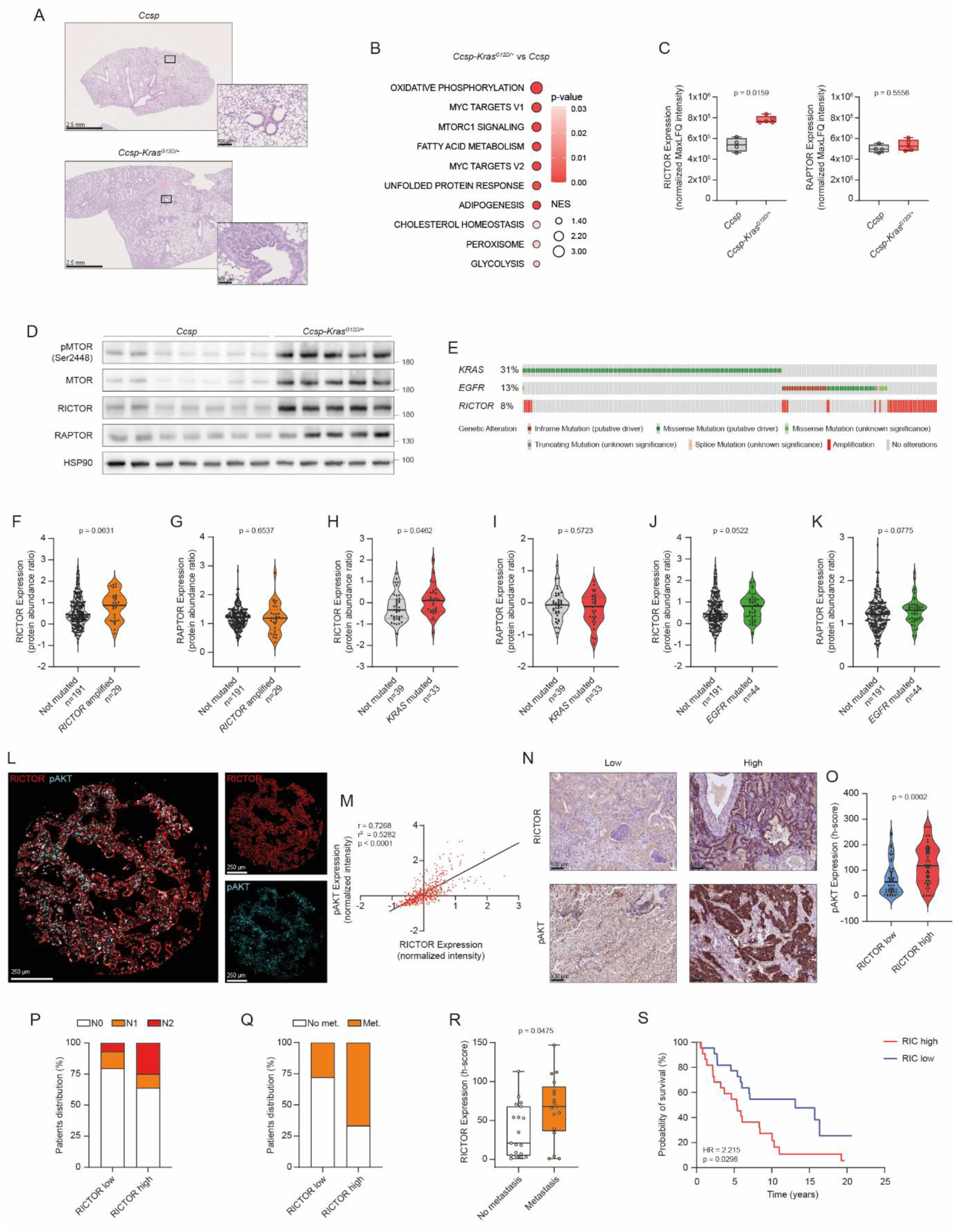
mTORC2 activation is a recurrent feature of aggressive lung tumors. **A.** Representative hematoxylin and eosin (H&E)-stained lung sections from *Ccsp* and *Ccsp-Kras^G12D/+^* mice. Scale bar represents 2.5 mm (left) and 100 µm (right). **B.** Gene set enrichment analysis (GSEA) performed on the proteomic analysis of lungs isolated from *Ccsp-Kras^G12D/+^* (n = 5) and *Ccsp* mice (n = 4). Shown are the significantly enriched Hallmark pathways (p < 0.05) ranked by normalized enrichment score (NES) and colored by significance (p-value). **C.** Boxplot of normalized MaxLFQ intensities for RICTOR and RAPTOR representing protein abundance across *Ccsp* (n = 4) and *Ccsp-Kras^G12D/+^* (n = 5) mice (Mann-Whitney U test). **D.** Western blot (WB) analysis of phosphorylated-MTOR (pMTOR Ser2448), MTOR, RICTOR and RAPTOR protein expression in lungs isolated from *Ccsp* and *Ccsp-Kras^G12D/+^* mice. HSP90 was used as a sample loading control. Representative image from one experiment with 7 animals for *Ccsp* and 5 for *Ccsp-Kras^G12D/+^*^.^ **E.** Oncoprint representation of *KRAS* mutations, *EGFR* mutations and *RICTOR* amplification in TCGA lung adenocarcinoma (LUAD) patients, performed via www.cbioportal.org. **F, G.** Violin plots showing RICTOR (**F**) and RAPTOR (**G**) protein abundance in LUAD samples stratified by *RICTOR* amplification (TCGA dataset, n = 220, Mann-Whitney U test). **H, I.** Violin plots showing RICTOR (**H**) and RAPTOR (**I**) protein abundance in LUAD samples stratified by *KRAS* mutational status (CPTAC dataset, n = 72, Mann-Whitney U test). **J, K.** Violin plots showing RICTOR (**J**) and RAPTOR (**K**) protein abundance in LUAD samples stratified by *EGFR* mutational status (TCGA dataset, n = 235, Mann-Whitney U test). **L.** Representative multiplex immunofluorescence image of a lung cancer biopsy stained for RICTOR (red) and phosphorylated-AKT on Ser473 (pAKT, light blue). The merged image illustrates co-localization, with corresponding single-channel images for RICTOR and pAKT Ser473. Scale bar represents 250 µm (left and right). **M.** Scatter plot showing the correlation between RICTOR and pAKT Ser473 protein levels, as quantified by multiplex immunofluorescence in lung cancer patient samples (Glasgow cohort, n = 463, Pearson correlation, Tumor Region only). **N.** Representative immunohistochemistry (IHC) images of lung cancer biopsies displaying low and high RICTOR and pAKT Ser473 expressions. Scale bar represents 100 µm. **O.** Violin plot showing pAKT Ser473 protein abundance (expressed as histoscore (h-score)) in lung cancer biopsies stratified by RICTOR protein expression (Cologne cohort, n = 57 for RIC^low^ and n = 50 for RIC^high,^ Mann-Whitney U test). **P.** Bar plot showing the differential distribution of lymph node infiltration (N stage) among lung cancer patients stratified according to RICTOR expression (Cologne cohort, n = 44 for RIC^low^ and n = 36 for RIC^high)^. **Q.** Bar plot showing the differential distribution of distant metastasis (met.) among lung cancer patients stratified according to RICTOR expression (Liège cohort, n = 18 for RIC^low^ and n = 18 for RIC^high)^. **R.** Boxplot of RICTOR protein abundance in non-metastatic and metastatic lung cancer patients biopsied before diagnostic (Liège cohort, n = 19 for non-metastatic and n = 17 for metastatic patients, median ± IQR, Mann-Whitney U test). **S.** Kaplan–Meier survival curves of lung cancer patients stratified according to RICTOR expression (Dataset obtained from Lehtiö et al (63), n = 22 for RIC^high^ (upper quartile) and n = 22 for RIC^low^ (lower quartile), hazard ratio (HR) determined by log-rank (Mantel–Cox) test).\

## mTORC2 activity promotes lung tumor progression

To directly interrogate the functional contribution of mTORC2 signaling to lung tumor progression, we sought to genetically inactivate *RICTOR*, the defining component of the mTORC2 complex. In agreement with previous studies (13), *RICTOR* silencing in human lung cancer cell lines impaired long-term clonogenic survival, triggered cell death and markedly reduced tumor engraftment in immune-deficient mice (Fig. S2A-C). To evaluate the physiological relevance and the therapeutic potential of mTORC2 inhibition in lung cancer, we generated genetically engineered mouse models (GEMMs) enabling conditional *Rictor* inactivation in *Kras*-driven lung tumors, either alone or in combination with *Trp53* loss (generating *Kras^G12D/+^*, *Kras^G12D/+;^Rictor^lox/lox^*, *Kras^G12D/+;^Trp53^lox/lox^* and *Kras^G12D/+;^Trp53^lox/lox;^Rictor^lox/lox^* mice, further referred to as K, KR_KO_, KP and KPR_KO_ mice respectively; Fig. 2A-E). Lung-specific recombination was induced by intratracheal administration of lung-tropic Cre-containing Adeno-Associated Viruses (AAV6.2ff Spb-Cre). Strikingly, genetic ablation of *Rictor*, hence mTORC2 inactivation, significantly reduced tumor burden in both KR_KO_ and KPR_KO_ models, when compared to their respective controls (Fig. 2A-D). In the KP model, *Rictor* deletion also markedly prolonged mouse survival (Fig. 2E). Importantly, conditional deletion of *Rictor* in normal lung epithelium did not induce detectable histological abnormalities several months after recombination (Fig. S2D), suggesting that mTORC2 activity is dispensable for normal lung homeostasis but selectively required for tumor progression.

**Figure 2:**
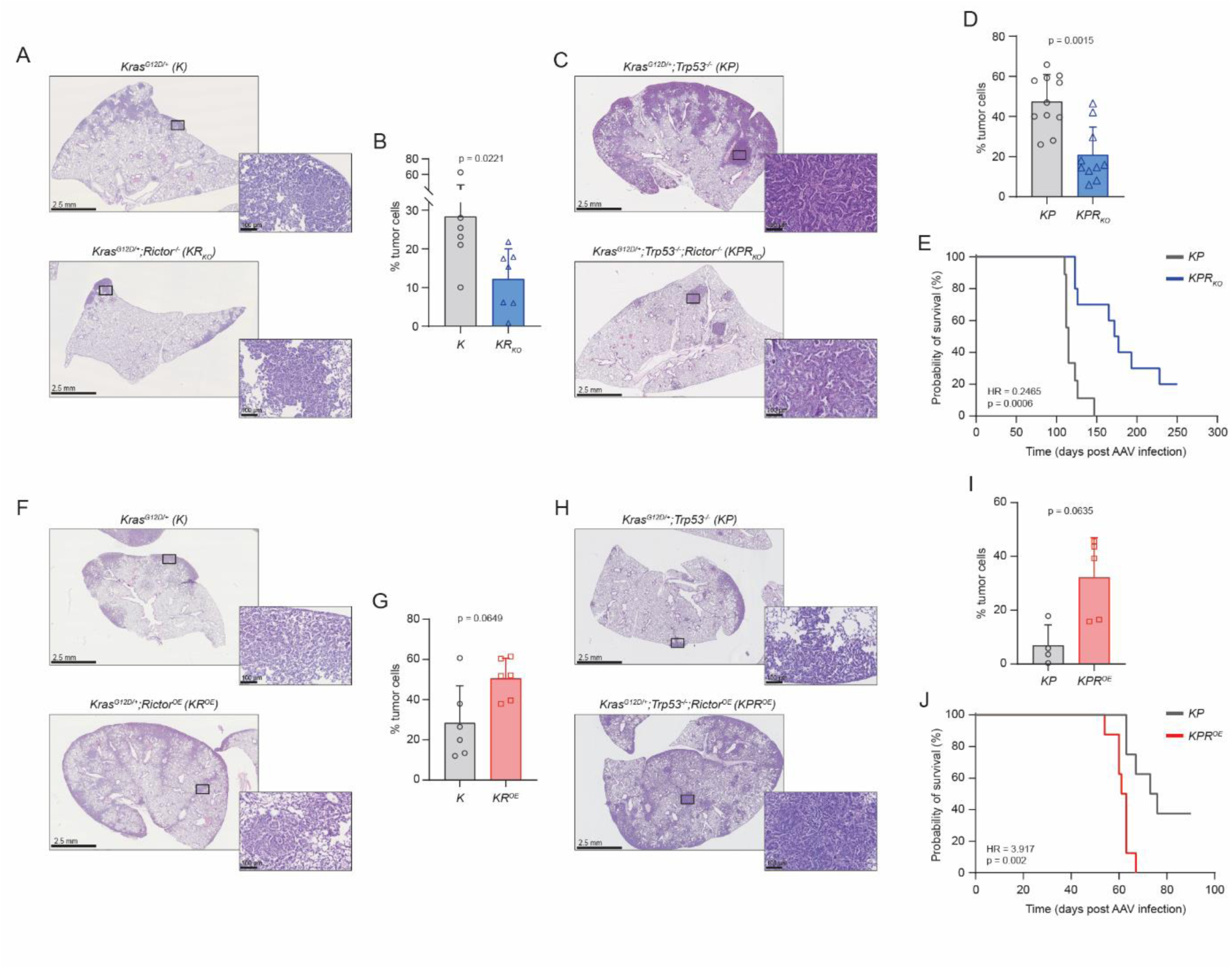
mTORC2 activity dictates lung tumor aggressiveness. **A.** Representative H&E-stained lung sections from *Kras^G12D/+^* (K) and *Kras^G12D/+;^Rictor^-/-^* (KR_KO_) mice. Scale bar represents 2.5 mm (left) and 100 µm (right). **B.** Quantification of tumor cell percentage in lungs from K (n = 6) and KR_KO_ (n = 7) mice (mean ± SD, Mann-Whitney U test). **C.** Representative H&E-stained lung sections from *Kras^G12D/+;^Trp53^-/-^*^(^KP) and *Kras^G12D/+;^Trp53^-/-;^Rictor^-/-^* (KPR_KO_) mice. Scale bar represents 2.5 mm (left) and 100 µm (right). **D.** Quantification of tumor cell percentage in lungs from KP (n = 11) and KPR_KO_ (n = 10) mice (mean ± SD, Mann-Whitney U test). **E.** Kaplan–Meier survival curves of KP (n = 9) and KPR_KO_ (n = 10) mice (HR determined by log-rank (Mantel-Cox) test). **F.** Representative H&E-stained lung sections from *Kras^G12D/+^* (K) and *Kras^G12D/+;^Rictor^OE^* (KR^OE)^ mice. Scale bar represents 2.5 mm (left) and 100 µm (right). **G.** Quantification of tumor cell percentage in lungs from K (n = 6) and KR^OE^ (n = 6) mice (mean ± SD, Mann-Whitney U test). **H.** Representative H&E–stained lung sections showing tumors from *Kras^G12D/+;^Trp53^-/-^* (KP) and *Kras^G12D/+;^Trp53^-/-;^Rictor^OE^* (KPR^OE)^ mice. Scale bar represents 2.5 mm (left) and 100 µm (right). **I.** Quantification of tumor cell percentage in lungs from KP (n = 4) and KPR^OE^ (n = 5) mice (mean ± SD, Mann-Whitney U test). **J.** Kaplan–Meier survival curves of KP (n = 8) and KPR^OE^ (n = 8) mice (HR determined by log-rank (Mantel-Cox) test).

Next, we sought to determine whether sustained mTORC2 activation could potentiate lung tumor development. To model this clinically relevant setting, we generated complementary GEMMs allowing constitutive or AAV-inducible overexpression of *Rictor* in the lung epithelium (*Ccsp-Cre;Rictor^OE^*, further referred to as *Ccsp-Rictor^OE^*, or *Rictor^OE^* models, respectively). *Rictor* overexpression alone did not alter lung architecture and was insufficient to induce tumorigenesis (Fig. S2E-F). However, sustained mTORC2 activation in *Kras*-driven lung tumors (*Kras^G12D/+;^Rictor^OE^* and *Kras^G12D/+;^Trp53^lox/lox;^Rictor^OE^* mice, further referred to as KR^OE^ and KPR^OE)^ significantly increased tumor burden (Fig. 2F-I) and reduced overall survival (Fig. 2J), consistent with patient data (Fig. 1S). Together, these findings identify mTORC2 signaling as a key contributor of lung cancer progression and support mTORC2 targeting as a promising strategy in lung cancer.

## mTORC2 activation drives the metabolic rewiring of lung tumors

Despite the strong therapeutic rationale for targeting mTORC2 signaling in cancer, the development of selective mTORC2 inhibitors remains challenging. We therefore reasoned that identifying dependencies downstream of mTORC2 could reveal therapeutically actionable vulnerabilities in lung cancer. To this end, we performed unbiased proteomic analyses of macro-dissected murine lung tumors following mTORC2 activation or inhibition (KPR^OE^ vs. KP; KPR_KO_ vs. KP). These datasets were further integrated with publicly available proteomic profiles of LUAD patients stratified according to intratumoral RICTOR levels (RIC^high^ vs. RIC^low^ tumors). GSEA identified consistent enrichment of pathways associated with cell proliferation and metabolic regulation in mTORC2-active tumors across both murine and human datasets (enriched in RIC^high^ and KPR^OE^ but underrepresented in KPR_KO_ tumors when compared to respective controls, Fig. 3A-C). By contrast, inflammatory and interferon-related pathways, were negatively associated with mTORC2 activation (Fig. S3A-B). Notably, glycolysis emerged as the metabolic program most consistently linked to mTORC2 activity across all three models (Fig. 3A-C). Consistent with this observation, western blot analyses confirmed altered expression of key glycolytic enzymes upon modulation of RICTOR expression in murine lung tumors (Fig. S3C-D). Furthermore, metabolomic profiling of RICTOR-depleted A549 lung cancer cells revealed reduced abundance of glucose-derived metabolites following mTORC2 inhibition (Fig. 3D).

**Figure 3:**
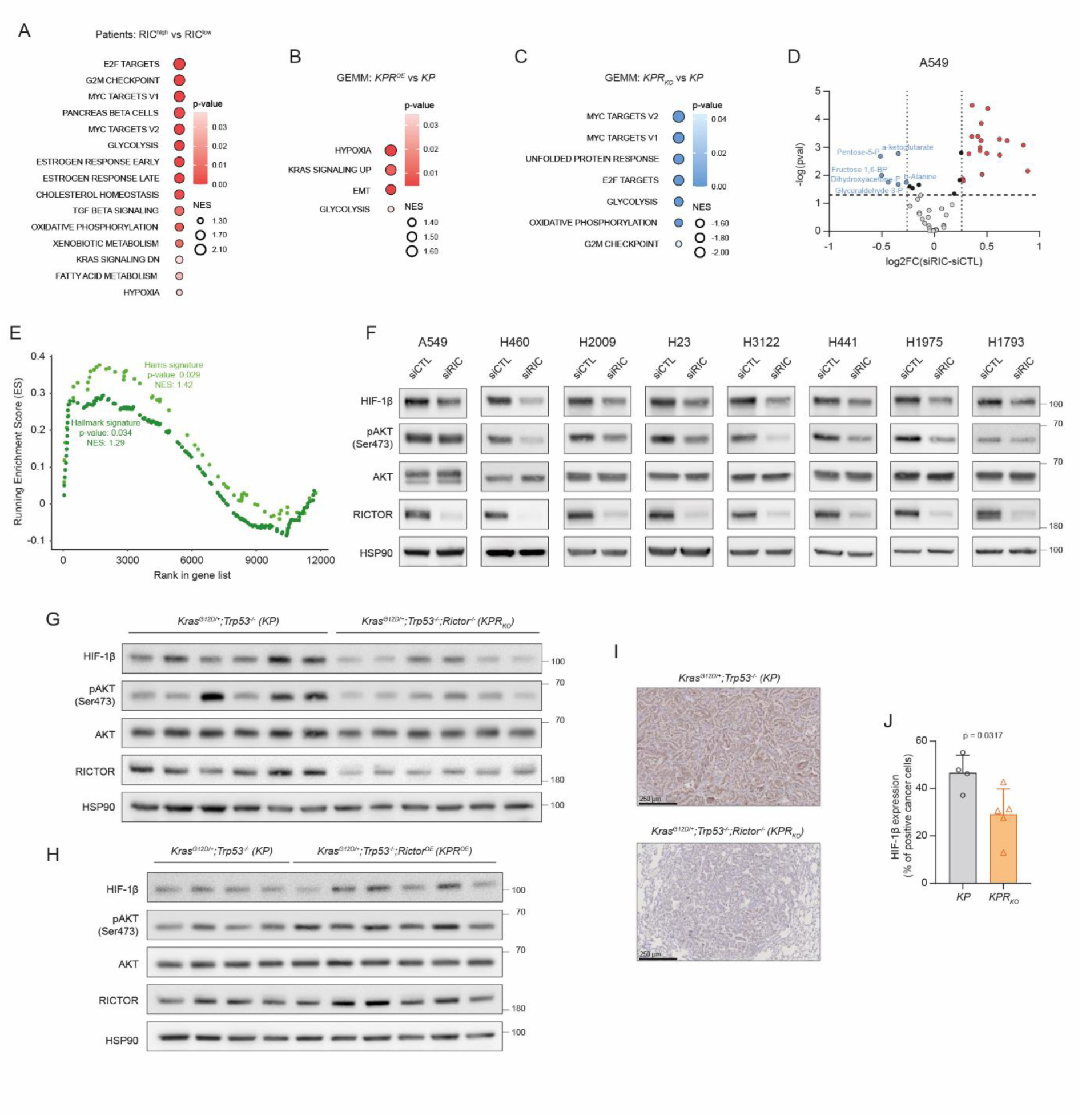
mTORC2 activation reprograms lung tumor metabolism. **A.** GSEA performed on proteomics of lung cancer biopsies from RICTOR high patients (RIC^high,^ n = 22, upper quartile) compared to RICTOR low patients (RIC^low,^ n = 22, lower quartile). Data was retrieved from Lehtiö et al (63). Shown are the significantly enriched Hallmark pathways ranked by NES and colored by significance (p-value). **B, C.** GSEA performed on proteomics of macro-dissected lung tumors isolated from KPR^OE^ mice (n = 6) (**B**) and KPR_KO_ mice (n = 5) (**C**) compared to their respective controls (n = 4 and n = 6, respectively). Shown are the significantly enriched (**B**) or downregulated (**C**) Hallmark pathways ranked by NES and colored by significance (p-value). **D.** Volcano plot of the differentially abundant metabolites in RICTOR-depleted (siRIC) and control (siCTL) A549 cells. Metabolites with significantly increased or decreased levels (p-value < 0.05) are highlighted in red and blue, respectively (n = 3 independent biological experiments). **E.** Enrichment score of the Hypoxia Hallmark signature and the Hypoxia Harris signature (64) applied on the proteomic analysis of lung cancer biopsies obtained from Lehtiö et al (63). **F.** WB analysis of HIF-1β, pAKT Ser473, AKT and RICTOR protein expression following RICTOR silencing in multiple human lung cancer cell lines. HSP90 was used as a sample loading control. Representative image from three independent biological experiments. **G.** WB analysis of HIF-1β, pAKT Ser473, AKT and RICTOR protein expression in macro-dissected lung tumors isolated from KP and KPR_KO_ mice. HSP90 was used as a sample loading control. Representative image from one experiment with 6 animals per group. **H.** WB analysis of HIF-1β, pAKT Ser473, AKT and RICTOR protein expression in macro-dissected lung tumors from KP and KPR^OE^ mice. HSP90 was used as a sample loading control. Representative image from one experiment with 4 animals for KP and 6 for KPR^OE.^ **I.** Representative IHC staining of HIF-1β expression in KP and KPR_KO_ mice. Scale bar represents 250 µm. **J.** Quantification of HIF-1β staining in tumor regions from KP (n = 4) and KPR_KO_ (n = 5) lungs (mean ± SD; Mann-Whitney U test).

### HIF-1β is a conserved and functional effector of mTORC2 signaling in lung cancer

Hypoxia is a major regulator of glycolysis (28). Interestingly, hypoxia signaling was significantly enriched in mTORC2-activated tumors (mouse and human, Fig. 3A-B) and was conversely down-regulated in RICTOR-depleted cells (Fig. S3E). Although mTORC2 signaling has previously been implicated in the regulation of glucose metabolism in cancer (29), hypoxia responses are classically considered to be predominantly controlled by mTORC1 activity (30,31). These observations prompted us to evaluate a potential role for mTORC2 in hypoxia signaling in lung cancer. Analyses of human LUAD datasets confirmed significant enrichment of hypoxia-related gene signatures in RIC^high^ tumors (Fig. 3E). However, neither hypoxia induction nor HIF-1α chemical stabilization activated mTORC2 signaling in lung cancer cells (Fig. S4A-B), suggesting that enhanced hypoxia signaling in RIC^high^ lung tumors arises as a consequence of increased mTORC2 activity rather than acting upstream of mTORC2. We therefore examined the expression of the hypoxia-inducible factors (HIF-1α, HIF-1β and HIF-2α) following transient RICTOR silencing in a panel of lung cancer cell lines. Strikingly, HIF-1β protein levels were consistently reduced upon RICTOR depletion (Fig. 3F). Moreover, HIF-1β expression strongly correlated with RICTOR abundance in stably-transduced cells (Fig. S4C), tumor xenografts (Fig. S4D) and, most importantly, in GEMM-derived lung tumors (Fig. 3G-J). Notably, this association was specific to HIF-1β, as the levels of HIF-1α and HIF-2α displayed heterogeneous responses to mTORC2 modulation across models (Fig. S4D-G). This differential regulation was particularly evident in RICTOR-depleted A549 xenografts, which exhibited reduced levels of HIF-1β together with increased HIF-1α and HIF-2α expression (Fig. S4D). Collectively, these results highlight HIF-1β as a major downstream effector of mTORC2 signaling in lung cancer and support a previously underappreciated role for HIF-1β in metabolic adaptation.

## A non-canonical mTORC2-PKCα-CK2 axis controls HIF-1β protein stability

We next sought to elucidate the molecular mechanisms linking mTORC2 activity to HIF-1β regulation. Pharmacological inhibition of mTOR signaling using the dual mTOR inhibitor PP-242 markedly decreased HIF-1β expression (Fig. 4A, Fig. S5A). Similarly, RICTOR depletion, but not RAPTOR silencing, reduced HIF-1β levels, confirming a selective role for mTORC2 in HIF-1β regulation (Fig. 4B). Unexpectedly, pharmacological inhibition of AKT, the canonical downstream effector of mTORC2, using MK-2206 failed to alter HIF-1β expression (Fig. 4C, Fig. S5B), suggesting the involvement of a non-canonical signaling pathway. Protein kinase C alpha (PKCα) is another well-established substrate for mTORC2 (11). Moreover, PKC has been reported to regulate casein kinase 2 (CK2) activity (32), which can in turn modulate HIF signaling (33). Consistent with this model, pharmacological inhibition of PKCα (using the pan-PKC inhibitor sotrastaurin) or CK2 (using silmitasertib) significantly reduced HIF-1β levels (Fig. 4D-E, Fig. S5C-D) in A549 and H460 lung cancer cells. Likewise, genetic silencing of PKCα or CK2α independently decreased HIF-1β levels in both cell lines, recapitulating the effects of pharmacological inhibition (Fig. 4F). Together, these findings support the existence of an AKT-independent mTORC2-PKCα-CK2 axis involved in the regulation of HIF-1β (Fig. 4G). Importantly, pharmacological inhibition of CK2 *in vivo* reduced tumor burden in the majority of KPR^OE^ mice (4 out of 5 mice, Fig. 4H-I), supporting the therapeutic relevance of this signaling node in lung cancer.

**Figure 4:**
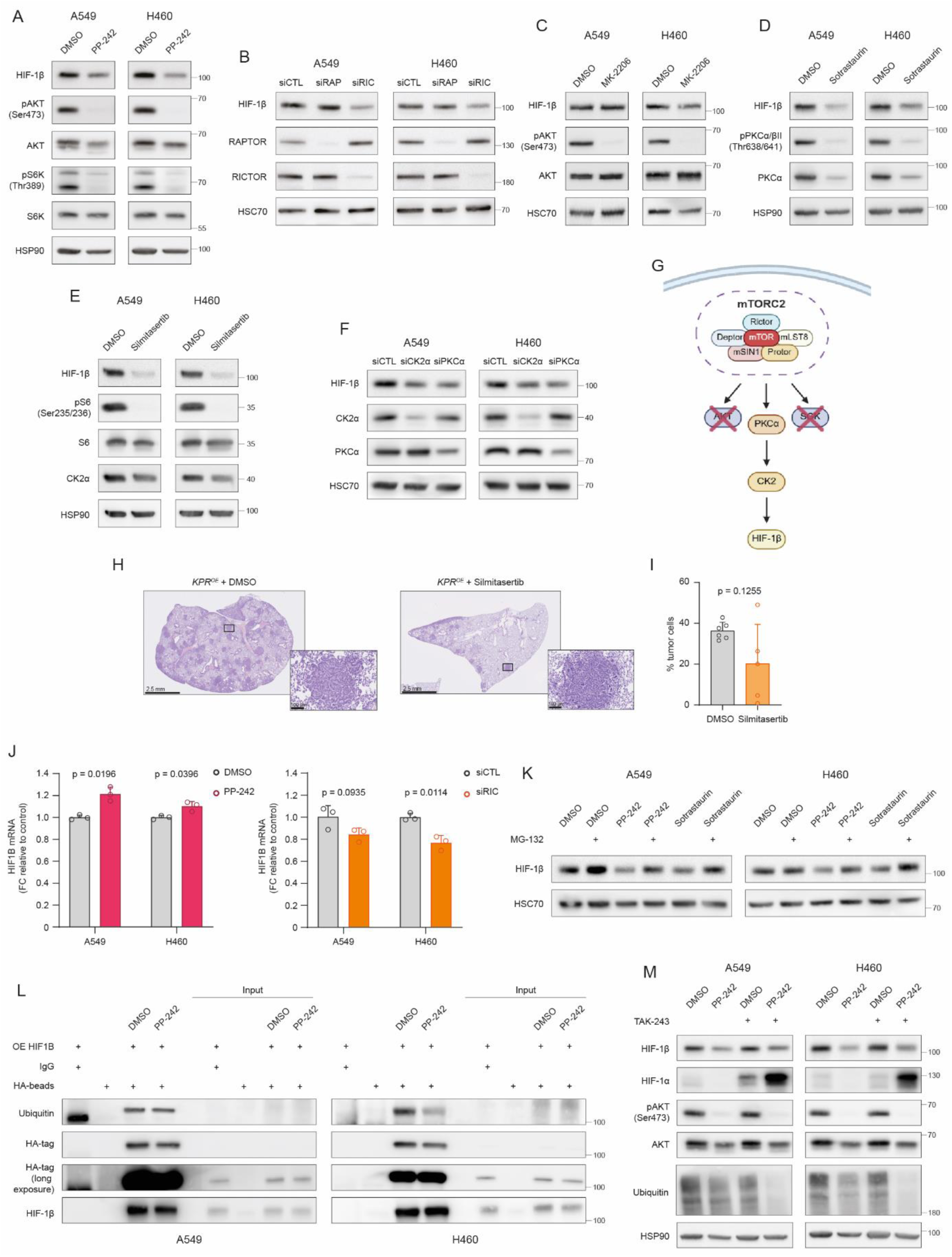
A non-canonical mTORC2-PKCa-CK2 axis controls HIF-1β protein stability. **A.** WB analysis of HIF-1β, pAKT Ser473, AKT, phosphorylated-P70S6K Thr389 (pS6K Thr389) and P70S6K (S6K) protein expression following 24h treatment with PP-242 (10µM) or vehicle control in A549 and H460 cell lines. HSP90 was used as a sample loading control. Representative image from three independent biological experiments. **B.** WB analysis of HIF-1β, RAPTOR and RICTOR protein expression following RICTOR and RAPTOR silencing in A549 and H460 cell lines. HSC70 was used as a sample loading control. Representative image from three independent biological experiments. **C.** WB analysis of HIF-1β, pAKT Ser473 and AKT protein expression following 24h treatment with MK-2206 (2µM) or vehicle control in A549 and H460 cell lines. HSC70 was used as a sample loading control. Representative image from three independent biological experiments. **D.** WB analysis of HIF-1β, phosphorylated-PKCα/β Thr638/641 (pPKCα/β Thr638/641) and PKCα protein expression following 24h treatment with sotrastaurin (50µM) or vehicle control in A549 and H460 cell lines. HSP90 was used as a sample loading control. Representative image from three independent biological experiments. **E.** WB analysis of HIF-1β, phosphorylated-S6 Ser235/236 (pS6 Ser235/236), S6 and CK2α protein expression following 24h treatment with silmitasertib (50µM) or vehicle control in A549 and H460 cell lines. HSP90 was used as a sample loading control. Representative image from three independent biological experiments. **F.** WB analysis of HIF-1β, CK2α and PKCα protein expression following CK2α and PKCα silencing in A549 and H460 cell lines. HSC70 was used as a sample loading control. Representative image from three independent biological experiments. **G.** Schematic representation illustrating the proposed signaling pathway regulating HIF-1β. Created with BioRender. **H.** Representative H&E– stained lung sections from KPR^OE^ mice treated with DMSO or silmitasertib. Scale bar represents 2.5 mm (left) and 100 µm (right). **I.** Quantification of tumor cell percentage in lungs from KPR^OE^ mice treated with DMSO (n = 6) or silmitasertib (n = 5) (mean ± SD, Mann-Whitney U test). **J**. Relative HIF1B mRNA expression in A549 and H460 cell lines following a 24h PP-242 (10µM) treatment (left) and RICTOR silencing (right), normalized to corresponding controls. Relative quantification of ΔΔCt is shown. (mean ± SD, n = 3 technical triplicates from a representative biological experiment (performed in independent biological triplicates), Welch’s t-test). GAPDH was used as a housekeeping gene. **K**. WB analysis of HIF-1β protein expression in A549 and H460 cell lines treated with DMSO, PP-242 (10µM) or sotrastaurin (50µM), in the presence or absence of MG-132 (20µM). HSC70 was used as a sample loading control. Representative image from three independent biological experiments. **L**. WB analysis of ubiquitin, HA-tag (short and long exposure) and HIF-1β following immunoprecipitation of HA-tagged HIF-1β in A549 and H460 cell lines treated with DMSO or PP-242 (10µM) for 8h in presence of MG-132 (20µM). Representative image from two (A549) or one (H460) independent biological experiment(s). **M**. WB analysis of HIF-1β, HIF-1α, pAKT Ser473, AKT and ubiquitin protein expression in A549 and H460 cell lines treated with DMSO or PP-242 (10µM), in the presence or absence of TAK-243 (1µM). HSP90 was used as a sample loading control. Representative image from three independent biological experiments.

Mechanistically, mTORC2 inhibition only moderately altered *HIF1B* mRNA expression (Fig. 4J) or translation of the *HIF1B* transcript, as assessed by polysome profiling (Fig. S5E-G). By contrast, proteasome inhibition using MG-132 restored HIF-1β levels following PP-242 and sotrastaurin treatments, indicating that mTORC2 signaling might regulate HIF-1β protein stability (Fig. 4K, Fig. S5H). Unexpectedly, immuno-precipitation experiments revealed that mTORC2 inhibition did not increase HIF-1β ubiquitination (Fig. 4L, Fig. S5I-J). This was in sharp contrast with HIF-1α, which became strongly poly-ubiquitinated under the same conditions (Fig. S5I-J). Similarly, global inhibition of ubiquitination using TAK-243, a selective ubiquitin activating enzyme (UBA1) inhibitor, failed to rescue HIF-1β degradation induced by PP-242 (Fig. 4M), demonstrating that, unlike HIF-1α, HIF-1β undergoes ubiquitin-independent proteasomal degradation upon mTORC2 inhibition.

Together, these findings identify a non-canonical mTORC2-PKCα-CK2 signaling axis that controls HIF-1β protein stability independently of AKT activity in lung cancer cells.

### HIF-1β links mTORC2 signaling to lipid metabolic reprogramming in lung cancer

Following the identification of HIF-1β as a downstream effector of mTORC2, we investigated the clinical and functional relevance of this signaling axis in lung cancer. Multiplex immunofluorescence imaging of human LUAD biopsies revealed a strong positive correlation between RICTOR and HIF-1β expression within tumor epithelial cells (r = 0.5576, Fig. 5A-B). Consistently, RIC^high^ tumors displayed significantly higher levels of HIF-1β than RIC^low^ lesions across two additional independent patient cohorts (Fig. 5C-E, r = 0.2934, Fig. S6A). This association was further preserved at the intratumoral level, as RIC^high^ regions were markedly enriched for HIF-1β compared to RIC^low^ areas within the same tumor (Fig. 5F-H). Importantly, concomitant high expression of RICTOR and HIF-1β identified a subgroup of patients characterized by increased metastatic dissemination and poor clinical outcome (36% of lymph node involvement and 75% of distant metastasis in RIC^highH^IF^high^ tumors versus 18.7% and 33.3% in RIC^lowH^IF^low^ tumors respectively, Fig. 5I-J), supporting a clinically relevant role for the mTORC2-HIF-1β axis in lung cancer progression.

**Figure 5:**
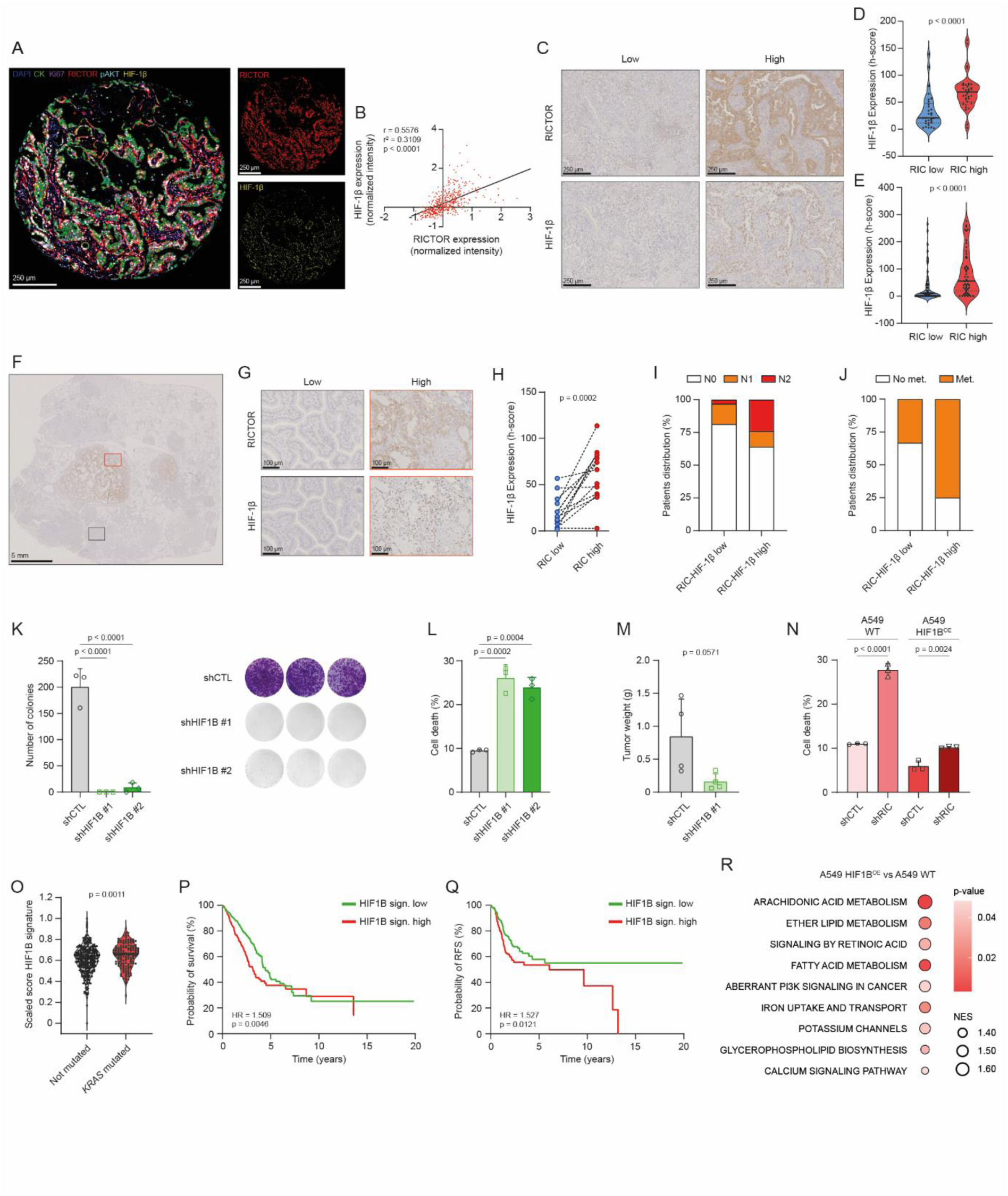
HIF-1β links mTORC2 signaling to lipid metabolic reprogramming in lung cancer. **A.** Representative multiplex immunofluorescence image of a lung cancer biopsy stained for DAPI (blue), Cytokeratin (CK, green), Ki67 (purple), RICTOR (red), pAKT Ser473 (light blue) and HIF-1β (yellow). The merged image illustrates co-localization or exclusion, with corresponding single-channel images for RICTOR and HIF-1β. Scale bar represents 250 µm (left and right). **B.** Scatter plot showing the correlation between RICTOR and HIF-1β protein abundance, as quantified by multiplex immunofluorescence in lung cancer biopsies (Glasgow cohort, n = 463, Pearson correlation, Tumor Region only). **C.** Representative IHC images of lung cancer patient biopsies displaying low and high expression of RICTOR and HIF-1β. Scale bar represents 250 µm. **D, E.** Violin plots showing HIF-1β expression levels (h-score) in lung cancer biopsies obtained from the Liège (**D**) and Cologne (**E**) cohorts, respectively, and stratified according to RICTOR expression (Liège cohort: n = 30 for RIC^low^ and n = 34 for RIC^high^ ; Cologne cohort: n = 57 for RIC^low^ and n = 50 for RIC^high,^ Mann-Whitney U test). **F, G.** Representative IHC images of a lung tumor biopsy displaying spatial heterogeneity in RICTOR and HIF-1β expression. Scale bar represents 5 mm (**F**) and 100 µm (**G**). **H**. Quantification of HIF-1β levels (h-score) in paired RIC^high^ versus RIC^low^ tumor regions (n = 14, Wilcoxon signed-rank test). **I.** Bar plot showing the distribution of lymph node infiltration (N Stage) among lung cancer patients stratified according to RICTOR and HIF-1β expression (Cologne cohort, n = 32 for RIC^lowH^IF^low^ and n = 25 for RIC^highH^IF^high)^. **J.** Bar plot showing the distribution of metastasis (met.) among lung cancer patients stratified according to RICTOR and HIF-1β expression (Liège cohort, n = 12 for RIC^lowH^IF^low^ and n = 12 for RIC^highH^IF^high)^. **K.** Quantification of colony formation assays performed with control (shCTL) and HIF1B-depleted (shHIF1B) A549 cells (mean ± SD, n = 3 technical triplicates from a representative biological experiment (performed in independent biological triplicates), one-way ANOVA with Dunnet’s correction test). **L.** Annexin V-PI staining (cell death) in shCTL and shHIF1B A549 cells (mean ± SD, n = 3 technical triplicates from a representative biological experiment (performed in independent biological triplicates), one-way ANOVA with Dunnet’s correction test). **M.** Endpoint tumor weights of shCTL and shHIF1B A549 xenografts grown in immunodeficient mice (mean ± SD, n = 4 tumors per group, Mann-Whitney U test). **N.** Annexin V-PI staining (cell death) in WT or HIF1B-overexpressing A549 cells (HIF1B^OE)^ depleted or not for RICTOR (shCTL and shRIC) (mean ± SD, n = 3 technical triplicates from a representative biological experiment (performed in independent biological triplicates), one-way ANOVA with Tukey’s correction test). **O.** Enrichment score of the HIF1B signature (top 100 up-regulated genes retrieved from the transcriptomics of HIF1B^OE^ A549 cell line) in non-mutated versus *KRAS*-mutated lung tumors (TCGA dataset, n = 301 for not mutated and n = 150 for *KRAS*-mutated tumors, Mann-Whitney U test). **P, Q.** Kaplan–Meier analysis of overall survival (OS) (**P**) and relapse-free survival (RFS) (**Q**) of lung cancer patients stratified according to the expression of the HIF1B signature (TCGA dataset, n = 265 (OS) and n = 233 (RFS) for HIF1B sign. low tumors, n = 248 (OS) and n = 212 (RFS) for HIF1B sign. high tumors, HR determined by log-rank (Mantel–Cox) test). **R.** GSEA performed on transcriptomic analysis of HIF1B^OE^ cells. Shown are the significantly enriched KEGG and Reactome pathways involved in metabolism, ranked by NES and colored by significance (p-value).

To directly assess the functional contribution of HIF-1β to lung tumorigenesis, we silenced its expression in lung cancer cells. HIF-1β depletion abrogated clonogenic growth, induced cell death and strongly reduced xenograft tumor growth *in vivo* (Fig. 5K-M, Fig. S6B), thereby phenocopying the effects observed following RICTOR depletion (Fig. S2A-C). Conversely, enforced HIF-1β overexpression rescued the survival defects induced by RICTOR silencing (Fig. 5N, Fig. S6C), demonstrating that HIF-1β functionally mediates key downstream effects of mTORC2 signaling in lung cancer cells. To further characterize HIF-1β-dependent transcriptional programs, we performed RNA sequencing analysis in HIF-1β-overexpressing A549 cells (HIF1B^OE^ cells). The resulting HIF1B gene signature (top 100 up-regulated genes in HIF1B^OE^ cells) was significantly enriched in *KRAS*-mutant lung tumors and associated with reduced overall and relapse-free survival in LUAD patients (Fig. 5O-Q). Notably, GSEA identified lipid metabolic pathways among the most significantly enriched programs downstream of HIF-1β (Fig. 5R), prompting us to investigate whether the mTORC2–HIF-1β axis directly controls lipid homeostasis in lung cancer.

## mTORC2-HIF-1β signaling creates a metabolic vulnerability in lung cancer

mTORC2 activation has previously been linked to lipid dysregulation in cancer (21). Consistently, RIC^high^ lung tumors displayed enrichment of lipid metabolic signatures (Fig. 3A). Because lipid metabolism emerged as a major transcriptional output of HIF-1β signaling, we investigated whether the mTORC2-HIF-1β axis directly regulates lipid homeostasis in lung cancer cells. Lipidomic profiling of A549 cells depleted of either RICTOR or HIF-1β (shRIC and shHIF1B cells, respectively) revealed extensive lipid remodelling following disruption of this pathway (Fig. 6A, Fig. S7A). In particular, multiple sphingolipid species were consistently under-represented in both RICTOR-and HIF-1β-deficient cells relative to their respective controls (Fig. 6A-C, Fig. S7A). Similar alterations were observed following transient siRNA-mediated silencing (Fig. S7B-C), suggesting a role for the mTORC2-HIF-1β axis in sphingolipid regulation.

**Figure 6:**
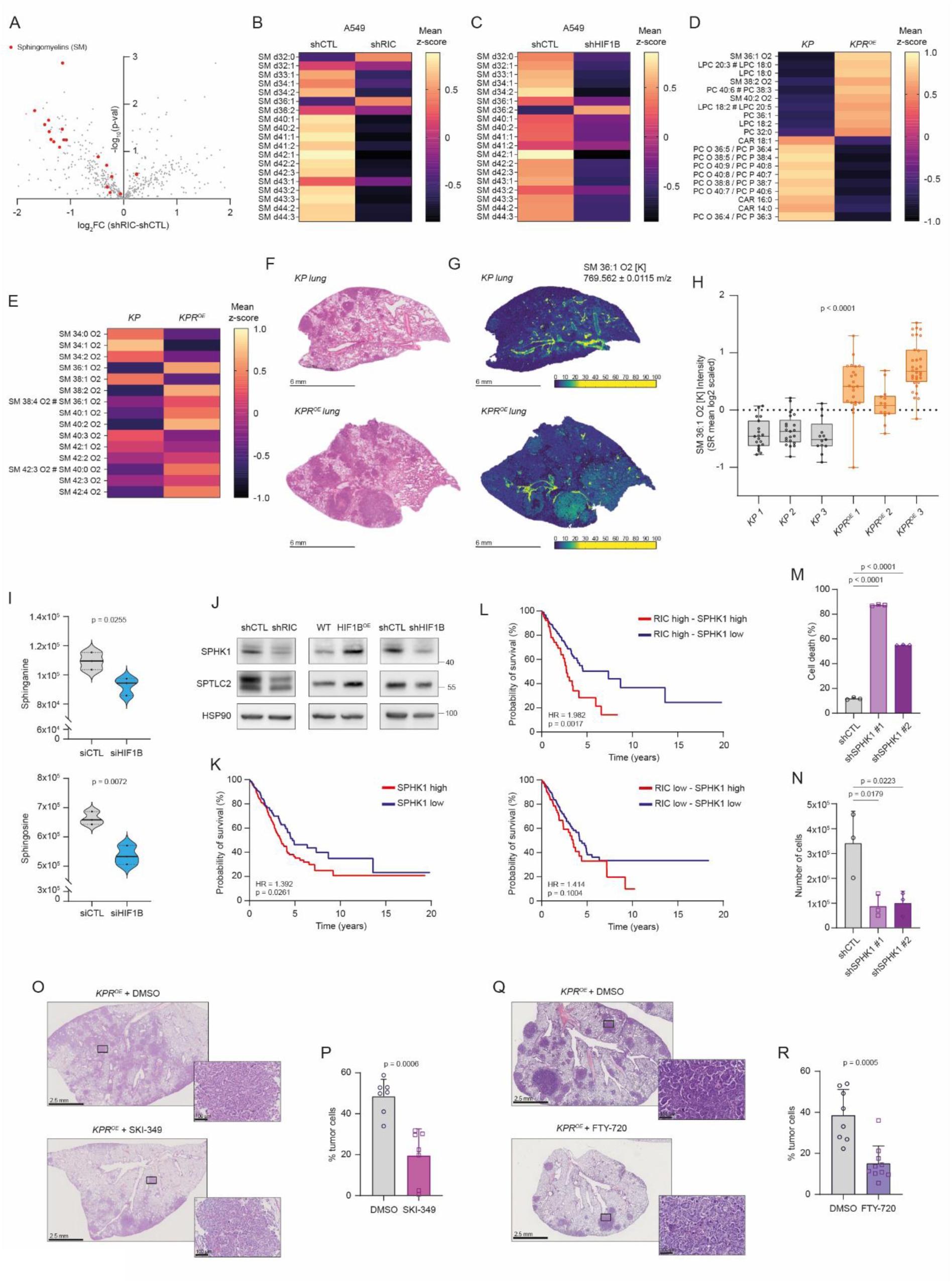
mTORC2-HIF-1β signaling creates a sphingolipid vulnerability in lung cancer. **A.** Volcano plot of the differentially abundant lipids in shRIC versus shCTL A549 cells. Sphingomyelins are highlighted in red (n = 3 independent biological experiments, Student t-test). **B, C.** Heatmaps illustrating the differential abundance of sphingomyelin species in A549 cells following RICTOR (shRIC) (**B**) or HIF1B (shHIF1B) (**C**) depletion (mean Z-scores, n = 3 independent biological experiments per group). **D.** Heatmap highlighting top 10 over-and under-represented lipids identified from spatial lipidomic analysis of KP and KPR^OE^ lung tumors (mean Z-scores, n = 3 animals per group). **E.** Heatmap illustrating the differential abundance of sphingomyelin species in KP and KPR^OE^ lung tumors (mean Z-scores, n = 3 animals per group). **F.** Representative H&E–stained lung sections from KP and KPR^OE^ mice. Scale bar represents 6 mm. Representative image from three independent animals per group. **G.** Representative image showing the spatial distribution and relative abundance of sphingomyelin SM 36:1 O2 in lung sections from KP and KPR^OE^ mice. Representative image from three animals per group. Scale bar represents 6 mm and color scale indicates relative intensity. **H.** Box plot displaying the normalized abundance of SM 36:1 O2 in lung tumor regions of KP and KPR^OE^ mice (n = 3 animals per group, False Discovery Rate with Benjamini-Hochberg correction). **I.** Violin plot showing sphinganine (top) and sphingosine (bottom) abundance in HIF1B-depleted A549 cells (siHIF1B) (n = 3 independent biological experiments for each group, Welch’s t-test). **J.** WB analysis of SPHK1 and SPTLC2 protein expression in shRIC (left), shHIF1B (right) and HIF1B^OE^ (middle) A549 cells. HSP90 was used as a sample loading control. Representative image from three independent biological experiments. **K.** Kaplan–Meier survival curves of lung cancer patients stratified according to SPHK1 expression (TCGA dataset, n = 245 for SPHK1^low^ tumors and n = 272 for SPHK1^high^ tumors, HR determined by log-rank (Mantel–Cox) test). **L.** Kaplan–Meier survival curves of lung cancer patients stratified according to RICTOR and SPHK1 expression (TCGA dataset, n = 67 for RIC^highS^PHK1^high^ tumors and n = 191 RIC^highS^PHK1^low^ tumors (upper panel), n = 71 for RIC^lowS^PHK1^high^ tumors and n = 188 RIC^lowS^PHK1^low^ tumors (lower panel), HR determined by log-rank (Mantel–Cox) test). **M, N.** Annexin V-PI staining (cell death) (**M**) and cell number (**N**) in control and SPHK1-depleted (shSPHK1) A549 cells (mean ± SD, for Annexin V-PI staining: n = 3 technical triplicates from a representative biological experiment (performed in independent biological triplicates); for cell number: n = 3 independent biological experiments, one-way ANOVA with Dunnet’s correction test). **O, Q.** Representative H&E-stained lung sections from KPR^OE^ mice treated with SKI-349 (**O**) or FTY-720 (**Q**) and their vehicle control. Scale bar represents 2.5 mm (left) and 100 µm (right). **P, R.** Quantification of tumor cell percentage in lungs from KPR^OE^ mice treated with DMSO (n = 7) and SKI-349 (n = 7) (**P**) or DMSO (n = 8) and FTY-720 (n = 10) (**R**) (mean ± SD, Mann-Whitney U test).

To determine whether mTORC2 activation similarly impacts lipid metabolism *in vivo*, we performed spatial lipidomic analyses on GEMM-derived lung tumors (KPR^OE^ vs KP) (Fig. 6D-H). Compared with KP tumors, KPR^OE^ lesions showed higher signal intensities of multiple sphingomyelin (SM) and lysophospholipid (LPC-LPE) species, whereas KP tumors displayed relatively higher signals for ether phospholipids and carnitines (Fig. 6D). More broadly, mTORC2 activation (in KPR^OE^ tumors) increased sphingolipid abundance *in vivo*, including marked accumulation of specific SM species such as SM 36:1 (Fig. 6E-H). Interestingly, HIF-1β depletion significantly reduced intracellular levels of sphinganine and sphingosine, two key intermediates of the sphingolipid biosynthetic pathway (Fig. 6I). In parallel, we identified serine palmitoyltransferase 2 (SPTLC2) and sphingosine kinase 1 (SPHK1) as downstream effectors of the mTORC2-HIF-1β axis, as their expression decreased following RICTOR or HIF-1β silencing and increased upon HIF-1β overexpression (Fig. 6J). Importantly, elevated SPHK1 expression was associated with poor clinical outcome in LUAD patients (Fig. 6K). This effect was particularly pronounced in tumors displaying high mTORC2-activity, where concomitant elevation of RICTOR and SPHK1 expression defined a subgroup of patients with markedly reduced survival (Fig. 6L), suggesting that mTORC2 activation establishes a clinically relevant sphingolipid vulnerability. To directly evaluate the therapeutic relevance of this pathway, we genetically and pharmacologically targeted sphingolipid metabolism in lung cancer models. SPHK1 inhibition markedly impaired cancer cell proliferation, clonogenic growth, and induced cell death (Fig. 6M-N, Fig. S7D-F). Most importantly, pharmacological inhibition of sphingolipid signaling using either the SPHK1 inhibitor SKI-349 or the sphingosine-1-phosphate receptor (S1PR) modulator FTY720 significantly reduced tumor burden in KPR^OE^ mice (Fig. 6O-R). Together, these findings uncover a mTORC2-HIF-1β-dependent sphingolipid vulnerability that can be therapeutically exploited in lung cancer.

## Discussion

Lung tumors display extensive genetic heterogeneity that limits the efficacy of current therapeutic strategies and promotes adaptive resistance. Identifying signaling pathways recurrently engaged across genetically distinct tumors may therefore reveal shared and therapeutically actionable vulnerabilities. Here, we identify mTORC2 signaling as a convergent adaptive dependency in lung cancer and demonstrate that RICTOR-dependent mTORC2 activity promotes tumor progression through HIF-1β-dependent metabolic rewiring. Using complementary GEMMs, multi-omics analyses and human patient cohorts, we uncover HIF-1b as a previously unrecognized effector of mTORC2 signaling that orchestrates metabolic adaptation and establishes a therapeutically exploitable vulnerability in lung cancer.

Our analyses across murine models and patient datasets reveal that elevated RICTOR protein expression and active mTORC2 signaling are recurrent features of molecularly distinct lung tumors, including *KRAS-*mutant*, EGFR-*mutant, and *RICTOR-*amplified lesions. Although these genetic alterations rarely co-occur in lung cancer, they appear to converge functionally on enhanced mTORC2 activity, suggesting that mTORC2 may represent a shared adaptive signaling node downstream of diverse oncogenic drivers. While therapeutic efforts in NSCLC have historically focused on mTORC1, accumulating evidence suggests that mTORC2 fulfills fundamentally distinct functions in tumor biology (14). Whereas mTORC1 primarily orchestrates anabolic growth programs, mTORC2 has emerged as a critical mediator of tumor adaptation by sustaining survival and metabolic plasticity under stress conditions (14,17,18). This adaptive role may be particularly relevant upon mTORC1 inhibition, where compensatory activation of PI3K-AKT signaling can maintain mTORC2 activity (34,35). Consistent with this model, we found that *KRAS*-driven lung tumors displayed evidence of selective mTORC2 activation associated with AMPKα-mediated suppression of mTORC1 signaling. These observations suggest that mTORC2 targeting may represent a more effective strategy than selective mTORC1 inhibition in lung cancer. Importantly, elevated RICTOR expression and mTORC2 activation in patient biopsies strongly correlated with metastatic progression and poor survival, supporting a clinically relevant role for mTORC2 signaling in lung tumor aggressiveness.

Genetic ablation of *Rictor* has previously established a critical role for mTORC2 signaling across multiple tumor types, including liver, pancreas, skin, breast, prostate and brain cancers (21,36–40). However, despite the extensive use of GEMMs to model *KRAS*-driven lung tumorigenesis, functional interrogation of mTORC2 signaling in this context has remained remarkably limited. Our findings therefore extend previous observations obtained in cell line-based systems and tumor xenografts, and establish the physiological relevance of mTORC2 signaling in oncogene-driven lung cancer progression *in vivo*. Using complementary gain-and loss-of-function mouse models, we demonstrate that mTORC2 activity is dispensable for normal lung homeostasis but critically required for efficient lung tumor progression. Conditional deletion of *Rictor* significantly reduced tumor burden and prolonged survival in *Kras*-driven models, including tumors lacking *Trp53*, whereas sustained *Rictor* overexpression markedly accelerated tumor progression and shortened survival. Importantly, mTORC2 activation alone was not sufficient to initiate tumorigenesis, supporting a model in which mTORC2 primarily functions as an adaptive signaling amplifier rather than an oncogenic driver. Together, these results identify mTORC2 signaling as a key and actionable dependency in lung cancer.

Despite the strong therapeutic rationale for targeting mTORC2 signaling in cancer, the development of selective mTORC2 inhibitors remains challenging. Current dual mTORC1/2 inhibitors lack specificity and have shown limited clinical efficacy (41). While emerging mTORC2-selective inhibitors such as JR-AB2-011 have demonstrated activity in preclinical models, issues related to specificity and the lack of patient stratification strategy have thus far precluded clinical translation (15,42). These challenges highlight the need to identify targetable vulnerabilities downstream of mTORC2. Our integrated proteomic, metabolomic and lipidomic analyses revealed that mTORC2 activation drives extensive metabolic rewiring in lung tumors, with glycolytic and lipid metabolic programs emerging among the most consistently enriched pathways across both murine and human datasets. A similar role for mTORC2 in coordinating glucose and lipid metabolism has been reported in the liver through AKT-and SREBP-dependent mechanisms (16). In cancer, aberrant mTORC2 signaling has subsequently been linked to lipid reprogramming, tumor progression and poor clinical outcome in hepatocellular carcinoma (21). Together, these findings support a conserved role for mTORC2 in coupling oncogenic signaling to metabolic adaptation across cancer types.

In this context, the strong enrichment of hypoxia-associated transcriptional programs in mTORC2-active lung tumors led us to identify the hypoxia-inducible factor HIF-1β as a key downstream effector of mTORC2 controlling tumor metabolism. Mechanistically, mTORC2 stabilized HIF-1β through a non-canonical PKCα-CK2 signaling axis independently of AKT activity and prevented its ubiquitin-independent proteasomal degradation. Further studies will be required to define the molecular determinants controlling HIF-1β degradation. Nonetheless, our findings reveal a distinct mechanism through which mTORC2 engages HIF-dependent adaptive programs. Functionally, HIF-1β depletion phenocopied the effects of RICTOR loss and markedly impaired lung tumor progression, whereas enforced HIF-1β expression rescued survival defects induced by mTORC2 inhibition. Taken together, these findings establish HIF-1β as a conserved and functional effector of mTORC2 signaling in lung cancer.

Although HIF-1β (also known as ARNT) has historically been viewed as a passive transcriptional partner in hypoxia signaling, accumulating evidence suggests broader functions in metabolic regulation (43). Beyond its established role as an obligate binding partner for HIF-1α, HIF-1β interacts with multiple transcription factors, including HIF-2α and AHR, to regulate diverse transcriptional programs involved in cellular adaptation and metabolism (44,45). Supporting this concept, HIF-1β participates in transcriptional programs controlling fatty acid synthesis and membrane lipid remodelling through both HIF-and AHR-dependent mechanisms (46,47). Moreover, genetic deletion of *Hif1b* in murine adipose tissue results in reduced adipocyte size and altered glucose metabolism (48), while HIF-1β signaling in the skin has been linked to the regulation of sphingolipid metabolism and epidermal barrier function (49,50). Consistent with these observations, transcriptomic and lipidomic analyses identified lipid metabolic programs, and particularly sphingolipid metabolism, as major downstream outputs of the mTORC2-HIF-1β axis. RICTOR or HIF-1β depletion markedly reduced sphingolipid abundance, whereas mTORC2 activation promoted accumulation of multiple sphingomyelin species *in vivo*. Moreover, we identified key enzymes involved in sphingolipid biosynthesis, including SPHK1 and SPTLC2, as downstream targets of the mTORC2-HIF-1β axis. Together, these findings support a model in which HIF-1β functions as a metabolic effector downstream of mTORC2 and coordinates lipid metabolic programs beyond canonical hypoxia signaling.

Sphingolipids are increasingly recognized as critical regulators of cancer cell fitness through their ability to control membrane organization and intra-and extracellular signaling (23). In particular, the balance between pro-apoptotic ceramides and pro-survival sphingosine-1-phosphate (S1P) is frequently disrupted in cancer and contributes to tumor progression, metastatic dissemination and therapeutic resistance (51). Consistent with this concept, multiple enzymes involved in sphingolipid homeostasis, including sphingosine kinases (SPHK1/2), ceramidases and ceramide synthases, are recurrently dysregulated in NSCLC and have been linked to oncogenic pathways such as PI3K-AKT-mTOR signaling (24–26). Our findings further establish sphingolipid metabolism as a clinically relevant downstream vulnerability of the mTORC2-HIF-1β axis. Elevated SPHK1 expression was associated with poor outcome in lung cancer patients, particularly in tumors displaying high mTORC2 activity, supporting the existence of a clinically relevant sphingolipid vulnerability in this molecular context. Importantly, pharmacological disruption of sphingolipid signaling using either the SPHK1 inhibitor SKI-349 or the S1PR modulator FTY720 markedly reduced tumor burden in mTORC2-driven lung tumors, demonstrating that mTORC2-dependent metabolic rewiring creates actionable therapeutic vulnerabilities. Although both compounds have demonstrated antitumor activity in preclinical lung cancer models (52,53), our findings mechanistically position sphingolipid metabolism as a targetable downstream vulnerability of mTORC2 signaling in lung cancer.

Together, our study identifies mTORC2 as a central signaling hub integrating oncogenic and metabolic programs in lung cancer. Rather than functioning as a primary oncogenic driver, mTORC2 appears to amplify tumor progression by promoting non-canonical stabilization of HIF-1β, thereby coupling oncogenic signaling to metabolic adaptation. By identifying HIF-1β as a selective metabolic effector downstream of mTORC2, our findings establish a previously unrecognized signaling axis that defines therapeutically actionable metabolic vulnerabilities in molecularly distinct lung tumors.

## Methods

### Cell lines and culture conditions

A549 (ATCC CCL-185), NCI-H460 (ATCC HTB-177), NCI-H23 (ATCC CRL-5800), NCI-H2009 (ATCC CRL-5911), NCI-H3122, NCI-H441 (ATCC HTB-174), NCI-H1975 (ATCC CRL-5908), NCI-H1793 (ATCC CRL-5896) cells were cultured in complete RPMI-1640 medium (L0501-500, Biowest, Nuaillé, France) supplemented with 10% foetal bovine serum (FBS) (Gibco, Thermo Fisher Scientific, Waltham, Massachusetts, USA), 1% L-glutamine (Gibco, Thermo Fisher Scientific, Waltham, Massachusetts, USA) and 1% penicillin/streptomycin (Gibco, Thermo Fisher Scientific, Waltham, Massachusetts, USA). Lenti-X 293T cells were cultured in DMEM (L0106-500, Biowest, Nuaillé, France) containing 10% FBS, 1% L-glutamine and 1% penicillin-streptomycin. All cell lines were maintained in 5% CO_2_ at 37°C and routinely tested for mycoplasma. H3122 and Lenti-X 293T cells were kindly provided by Prof. Reinhard Büttner (University of Cologne, Germany) and Dr. Emmanuel Di Valentin (GIGA-Institute, University of Liège, Belgium) respectively. All other cell lines were obtained from ATCC.

## Drug treatments

Cells were seeded at 2 × 10^5^ cells/well in 6-well plates and allowed to attach for 24h before treatment. The following reagents were purchased from Selleck Chemicals (Houston, Texas, USA) and resuspended in DMSO: Torkinib/PP-242 (S2218), MK-2206 (S1078), Sotrastaurin/AEB07 (S2791), Silmitasertib/CX-4945 (S0707), MG132 (S2619), TAK-243/MLN7243 (S8341), and DMOG/dimethyloxalylglycine (S7483). SKI-349 (HY-162143) was purchased from MedChemExpress (Monmouth Junction, New Jersey, USA) and also resuspended in DMSO. For hypoxia experiments, cells were placed in a hypoxia chamber (1% O₂, 5% CO₂) (Whitley H35 Hypoxystation, Don Whitley Scientific Limited, West Yorkshire, UK) and incubated for 24 h or 48 h before protein extraction.

## siRNA-mediated transfections

70%-confluent cells were transfected using Lipofectamine RNAimax reagent (Invitrogen, Thermo Fisher Scientific, Waltham, Massachusetts, USA) according to the manufacturer’s protocol. Following initial transfection, culture medium was refreshed after 8 h and the cells were allowed to grow for another 48 h before lysis. All siRNAs were purchased from Dharmacon (Lafayette, Colorado, USA): siNon-Targeting Control (siNT) pool (D-001810-10-20), siRICTOR human pool (L-016984-00-0005), siRPTOR human pool (L-004107-00-0005), siCSNK2A1 human pool (L-003475-00-0005), siPKC*a* human pool (L-003523-00-0005) and siHIF1B/siARNT human pool (L-007207-00-0005).

## shRNA-mediated transductions and stable overexpression experiments

Lenti-X 293T cells were stably transfected with lentiviral construction (12µg), pPsx2 (12µg) and VSV-G (5µg) plasmids using TransIT®-LT1 Transfection Reagent (Mirus Bio, MIR 2360, Fisher Scientific, Waltham, Massachusetts, USA) to produce the lentiviruses. After 48 h, virus-containing supernatant was centrifuged and filtered (0.45µm PVDF filter; HVLP04700, Milipore, Darmstadt, Germany) to collect the viruses, mixed with polybrene (8µg/mL; TR-1003-G, Sigma Aldrich, St Louis, Michigan, USA) and deposited on recipient cells (A549). A second round of transduction was repeated after 24 h to maximize transduction efficiency. Recipient cells were then selected with puromycin (1µg/mL; ant-pr-1, InvivoGen, San Diego, California, USA) or hygromycin B Gold (100µg/mL; ant-hg-1, InvivoGen, San Diego, California, USA), for additional 72 h. shRNA constructions were purchased from VectorBuilder (Chicago, Illinois, USA) and HA-tagged HIF1B expression plasmid was custom-synthesized by VectorBuilder. All constructs used in this study are listed in Supplementary Table 1.

## RNA extraction and quantitative real-time PCR

Total RNA was extracted using the E.Z.N.A. Total RNA Kit I (Omega Bio-Tek company, Norcross, Georgia, USA) according to the manufacturer’s protocol and quantified with NanoDrop 1000 Spectrophotometer (Thermo Fisher Scientific, Waltham, Massachusetts, USA). Reverse transcription of the eluted RNA was performed using the Thermo Scientific™ RevertAid H Minus First Strand cDNA synthesis Kit (Thermo Fisher Scientific, Waltham, Massachusetts, USA), using 1µg of total RNA as input and oligo-dT strategy. PCR was then performed with the corresponding primers (see Supplementary Table 2) diluted in SYBRGreen on the Light Cycler 480 II (Roche, Basel, Switzerland). GAPDH was used as the housekeeping gene to normalize relative gene expression.

## Protein extraction and western blotting

Mice tissues (lungs, lung tumors or tumor xenografts) were pulverized in liquid nitrogen. Cultured cells were washed twice with PBS and subsequently scraped in extraction buffer. Proteins from cells or tissue powder were extracted in an 1% SDS lysis buffer (supplemented with Complete™ protease inhibitor (Cat# 04693116001, Roche, Basel, Switzerland) and phosphatase inhibitor cocktail (Cat# 04906837001, Roche, Basel, Switzerland) and quantified with the BCA protein assay kit (Cat# 23227, Thermo Fisher Scientific, Waltham, Massachusetts, USA). Equal amounts of protein (diluted in 5x Laemmli sample buffer) were loaded on acrylamide gels for the migration (SDS-PAGE) and transferred on PVDF membranes (IPVH85R, Merck Millipore, Burlington, Massachusetts, USA). Blocking was performed in 5% non-fat milk diluted in TBS-T (Tris-buffered saline with 0.1% Tween-20) for 1 h at room temperature. Membranes were then incubated at 4°C overnight with the primary antibody (see supplementary Table 3 for references and dilutions). After three washes with TBS-T, HRP-conjugated secondary antibody (anti-mouse or anti-rabbit) diluted in 5% non-fat milk was added for 1 h at room temperature. Membranes were then revealed using chemiluminescence (ECL, Cat# 32106, Thermo Fisher Scientific, Waltham, Massachusetts, USA) and signal was acquired on an ImageQuant LAS 4000 mini revealing device (GE Healthcare, Chicago, Illinois, USA). ImageJ (v1.54, National Institutes of Health, Bethesda, Maryland, USA) was used to generate the final files. HSP90 or HSC70 were used as sample loading controls.

## Colony formation assay

Following antibiotic selection, stably transduced cells (A549) were replated in 6-well plates at a concentration of 1× 10^3^ cells/mL (3 mL/well) in complete RPMI-1640 medium (L0501-500, Biowest, Nuaillé, France). The plates were then incubated for 7-14 days at 37°C. Once colonies were established, cells were gently washed with PBS, fixed with 4% paraformaldehyde for 15 min at room temperature and stained with 0.5% (w/v) Crystal Violet (diluted in 20% methanol) for 20 min. Images were acquired with a digital camera and colonies were counted with ImageJ (v1.54, National Institutes of Health, Bethesda, Maryland, USA).

## Cell death assay (FACS Annexin V/PI)

Adherent and floating cells were collected and stained with Annexin V-FITC and propidium iodide (PI) using the Dead Cell Apoptosis Kit for Flow Cytometry (V13242, Thermo Fisher Scientific, Waltham, Massachusetts, USA) according to the manufacturer’s instructions. Stained cells were analyzed by flow cytometry using FACSCanto II (BD Biosciences, Franklin Lakes, New Jersey, USA). Data were analyzed on FlowJo (v10.0.7, BD Biosciences, Franklin Lakes, New Jersey, USA) and dead cells were defined as a combination of Annexin V+/PI+, Annexin V^+/^PI^-a^nd Annexin V^-/^PI^+.^

## Immunoprecipitation

Cells were lysed in ice-cold lysis buffer (50mM TrisHCl (pH 8), 150mM NaCl, 1% NP40, 1 tablet of Complete™ protease inhibitor cocktail (Cat# 04693116001, Roche, Basel, Switzerland) for 50mL buffer). Protein content was determined using BCA protein assay kit (Cat# 23227, Thermo Fisher Scientific, Waltham, Massachusetts, USA). For the precipitation of ubiquitinated proteins, 1 mg of proteins per sample were incubated with Agarose-TUBE2 beads (Cat. #UM402, LifeSensors, Malvern, Pennsylvania, USA) or corresponding control agarose beads (Cat. #UM400, LifeSensors, Malvern, Pennsylvania, USA) for 1 h at 4°C. For the pull down experiments (HA-tagged HIF1B), 1 mg of proteins per sample were incubated with anti-HA-Tag agarose beads (sc-7392 AC, Santa Cruz Biotechnology, Dallas, Texas, USA) or control mouse IgG antibody (sc-2025, Santa Cruz Biotechnology, Dallas, Texas, USA) overnight at 4°C. Protein G PLUS-agarose beads (sc-2002, Santa Cruz Biotechnology, Dallas, Texas, USA) were then added to control IgG lysates for 1 h at 4°C. After incubation, complexes were extensively washed with lysis buffer and then eluted in 5x Laemmli sample buffer to be analyzed by western blot. Inputs were collected and ran on the same western blot.

## Murine xenograft tumor model

All animal procedures were performed in accordance with European ethical guidelines and were approved by the local animal care and use committee of the University of Liège (protocol license no. 20-2267, 20-2269 and 23-2559). Mice were housed the animal facility of the University of Liège and maintained under a light-dark cycle. Temperature was kept between 20-25°C. Food and water were provided ad libitum.

For xenograft experiments, 12 weeks-old NOD-SCID mice, grown at the facility, were randomly allocated into experimental groups at the time of injection. Cell injections were performed blindly by the investigator (not knowing which cell suspension was prepared for the injection). A549 cells were trypsinized, counted, washed twice with PBS and resuspended in serum-free RPMI-1640 medium (L0501-500, Biowest, Nuaillé, France) mixed 1:1 with Matrigel (Corning, New-York, USA) at a concentration of 3 × 10^6^ cells/200 µL/mouse. Tumor cells were then injected subcutaneously into the right flank of each mouse. At experimental endpoint, tumors were excised, measured and weighted. Half of the tumor material was fixed in 4% paraformaldehyde for subsequent histological studies and the other half was snap-frozen in liquid nitrogen and stored at -80°C for molecular analyses.

## Generation of mouse models

*R26-mRictor^tg^* mice were generated according to a previously described optimized targeting strategy (54). A recombinase-mediated cassette exchange (RMCE)-compatible targeting vector, *pRMCE-DV3-mRictor*, was constructed by a stepwise multisite Gateway LR reaction (Thermo Fisher Scientific, Waltham, Massachusetts, USA). Briefly, three entry vectors (pENTR L4R1 floxed transcriptional stop cassette, pENTR221 mRictor, pEntry 3’IRES-enhanced GFP (EGFP)/Luciferase) were combined overnight, followed by a second overnight LR reaction together with the *pRMCE-DV3* vector (54). Obtained pRMCE-DV3-mRictor clones were confirmed by restriction enzyme digests and sequencing analysis. The conditional cassette in the pRMCE-DV3-mRictor targeting vector was flanked by wild-type (FRTwt) and mutant (FRTmut) FRT sites and introduced into the *Rosa26* (R26) locus of G4 RosaLuc mouse embryonic stem cells (mESCs) by FLPe-mediated RMCE, as previously described (55). Correct integration into the *Rosa26* (R26) locus restored neomycin resistance (NeoR) expression, and was confirmed by genomic PCR. Correctly targeted mESC clones with a normal karyotype were aggregated with wild-type embryos and transferred into pseudopregnant females. Germline transmission was confirmed, leading to establishment of the *R26-mRictor^tg^* homozygous mouse line, referred to as *Rictor^OE^*^.^ Cre-mediated excision of the floxed transcriptional stop cassette activated expression of the bicistronic transgene transcript encoding for mRictor and an eGFP/Luciferase dual reporter, driven by the endogenous *Rosa26* promoter.

*LSL-Kras^G12D/+^* mice were obtained from The Jackson Laboratory (B6.129S4-Krastm4Tyj/J, The Jackson Laboratory, Bar Harbor, Maine, USA). *Trp53^lox/lox^* (*p53^lox/lox^*^)^ mice were a kind gift from Prof. Christophe Desmet (GIGA-Institute, University of Liège, Belgium). *Scgb1a1*-Cre mice (*Ccsp-Cre*) were provided by Prof. Thomas Marichal (GIGA-Institute, University of Liège, Belgium). *Rictor^lox/lox^* mice were obtained from Prof. Michael Hall (University of Basel, Switzerland) and described previously (56). The *Rictor^lox/lox^* mouse was crossed with the *LSL-Kras^G12D/+^* and the *Trp53*^l*ox/lox*^ strains to generate *Kras^G12D/+;^Rictor^lox/lox^* and *Kras^G12D/+;^p53^lox/lox;^Rictor^lox/lox^* mice. The *Rictor*^OE^ mouse was crossed with the *LSL-Kras^G12D/+^*, *Trp53*^l*ox/lox*^ and the *Scgb1a1*-Cre strains to generate *Kras^G12D/+;^Rictor^OE^*, *Kras^G12D/+;^p53^lox/lox;^Rictor^OE^* and *Ccsp-Cre;Rictor^OE^* mice respectively. For inducible models, Cre recombination was induced in 10-weeks old mice following intratracheal delivery of Cre-containing Adeno-Associated Viruses (AAV).

At endpoint, lungs were collected and either fixed in 4% neutral-buffered formalin (sc-281692, Santa Cruz Biotechnology, Dallas, Texas, USA) for 24 h and embedded into paraffin blocks for future HE or IHC staining, or freshly macro-dissected to excise tumors, snap-frozen in liquid nitrogen and stored at -80°C for future protein or RNA extractions.

## AAV vector production

Recombinant adeno-associated viral vectors (AAV) were produced by the GIGA-Viral Vector platform. To restrict the expression of Cre recombinase to lung epithelial cells, AAV6.2FF vectors (displaying high tropism toward alveolar epithelial cells (57) expressing the Cre recombinase under the specific promoter of surfactant protein B (#VB200603-1243gsg, VectorBuilder, Chicago, Illinois, USA) were generated. Briefly, pAAV mSP-B Cre IRES mCherry plasmids were co-transfected into 293AAV Cell Line (AAV-100, Cell Biolabs, San Diego, California, USA) together with a helper plasmid (Part No. 340202 VPK-401 kit, Cell Biolabs, San Diego, California, USA) and REP-Cap plasmid (pAAV 6.2FF, Cell Biolabs, San Diego, California, USA) and pAAV-2 (#VPK-422, Cell Biolabs, San Diego, California, USA). After collecting AAV from cells, rAAV vectors were concentrated and titrated using ABM good kit (#GE931, Applied Biological Materials, Richmond, British Columbia, Canada) at a concentration of >1E+13 genome copy/mL (GC/mL).

### *In vivo* drug treatment

*KPR^OE^* mice were randomized into control and treated groups at the start of the experiment. At early tumor onset (4 weeks post AAV infection), mice were treated i.p. with SKI-349 (15 mg/kg; HY-162143, MedChemExpress, Monmouth Junction, New Jersey, USA), Fingolimod/FTY-720 (5 mg/kg; S5950, Selleck Chemicals, Houston, Texas, USA) and Silmitasertib/CX-4945 (50 mg/kg; S0707, Selleck Chemicals, Houston, Texas, USA) or DMSO every 2 days for 14-21 days until reaching ethical endpoint. All compounds were resuspended in DMSO. Doses and routes of administration were chosen according to literature review.

## Patient material

For the Liege cohort, lung tumor biopsies were retrieved from the biobank of the University Hospital Center in Liege. Informed consent was obtained from all patients providing samples and the protocol was approved by the ethical committee of the University of Liege (no. 2022/68).

For the Cologne cohort, tissue samples were retrieved and used for immunohistochemistry from the biobank of the Medical Faculty, University Hospital Cologne. Sampling and usage of tissues was approved by the Ethics Committee, University of Cologne No. 13-091/2017.

The Glasgow (LATTICe) cohort was originally constructed under REC 14/EM/1159 (East Midlands REC), and the ongoing management of the resource by the Greater Glasgow and Clyde Biorepository was approved under an amendment granted by the Leicester South REC. Ongoing use of the collection is now managed under REC 16/WS/0207. Raw data, including images, relating to the LATTICe TMA cohort used in this publication are subject to NHSGGC Biorepository ethical approval. To access this data a biorepository application and MTA are required, please contact JLQ or the NHSGGC Biorepository manager for advice on the application process.

## Immunohistochemistry

For H&E staining, paraffin sections (5µm) were deparaffinized in xylene, rehydrated through graded ethanol and stained with hematoxylin (Sigma Aldrich, St Louis, Michigan, USA) and counterstained with eosin Y (Sigma Aldrich, St Louis, Michigan, USA) according to standard protocols. Slides were then dehydrated, cleared and mounted using DPX mounting medium (Sigma Aldrich, St Louis, Michigan, USA). Staining procedure was automated using the Leica ST5010 Autostainer XL (Equipment No.: 12610645, Leica Biosystems, Wetzlar, Germany). For IHC staining, paraffin sections were deparaffinized in xylene, rehydrated through graded ethanol, autoclaved for 11 min at 126°C in 10mM citrate buffer pH 6 (Dako Target Retrieval Solution, S2369, Agilent Technologies, Santa Clara, California, USA) and blocked with hydrogen peroxide (1.07209.0250, Merck KGaA, Darmstadt, Germany). Next, animal free blocking solution (#15019, Cell Signaling Technology, Danvers, Massachusetts, USA) was added before primary antibody incubation (HIF1β/ARNT (#5537, Cell Signaling Technology, Danvers, Massachusetts, USA): 1/200, 1 h at room temperature in Dako REAL™ Antibody Diluent (S2022, Agilent Technologies, Santa Clara, California, USA)). Finally, the HRP-conjugated secondary antibody (K4003, Agilent Technologies, Santa Clara, California, USA) was added before staining with SignalStain® DAB mix (#11724, Cell Signaling Technology, Danvers, Massachusetts, USA). Slides were then counterstained by the Leica ST5010 Autostainer XL (Equipment No.: 12610645, Leica Biosystems, Wetzlar, Germany) and mounted using the EUKITT (EUKITT®, Kindler Gmbh, Freiburg, Germany). Slides were scanned with an Hamamatsu NanoZoomer 2.0-HT slide scanner (Hamamatsu Photonics, Chuo-ku, Hamamatsu City, Japan). Tumor quantification was performed with QuPath software (v0.5.1, University of Edinburgh, Edinburgh, UK). Tumor and non-tumor regions were identified using a pixel classifier (mice samples) or object classifier (human samples). Tumor burden was calculated as a proportion of total tissue area (H&E staining). DAB-positive cells were identified and quantified in tumor regions (IHC staining). Whole lungs were always used for quantification to account for tumor heterogeneity.

## Multiplex Immunofluorescence Imaging

Twelve 4µm-thick formalin fixed paraffin embedded (FFPE) lung adenocarcinoma TMA samples were sectioned and placed on to TOMO hydrophilic adhesive microscope slides (Matsunami, Kishiwada City, Osaka, Japan) and baked in a 60℃ oven for a minimum of 30 min. Following antibody validation, automated multiplex immunofluorescent staining was performed on the Ventana Discovery Ultra platform (RUO Discovery Universal v21.00.0019, Roche Tissue Diagnostics, Oro Valley, Arizona, USA) with an Opal fluorophore tyramide-based signal amplification system (Akoya Biosystems, Marlborough, Massachusetts, USA). The multiplex was run with a baking step at 60℃, before a dewax step for 24 min at 69℃, followed by a CC1 (pH 9) pretreatment for antigen retrieval was applied for 32 min at 95℃. Inhibitor CM (760-4840, Roche Tissue Diagnostics, Oro Valley, Arizona, USA) was applied for 12 min to block any previous endogenous peroxidase. The primary-secondary-antibody pairs used in the multiplex panel alongside their assigned opal fluorophore were applied in a sequential manner, respectively, are as follows: HIF-1β (5537S, Cell Signaling Technology, Danvers, Massachusetts, USA; 1/10, 4 h incubation), UltraMap-antiRb HRP (760-4315, Roche Tissue Diagnostics, Oro Valley, Arizona, USA; 24 min incubation), Opal 520 (OP001001, Akoya Biosystems, Marlborough, Massachusetts, USA; 1/100, 8 min incubation); p-AKT (EP2109Y) (ab81283, Abcam, Cambridge, UK; 1/25, 2 h incubation), OmniMap-antiRb HRP (760-4311, Roche Tissue Diagnostics, Oro Valley, Arizona, USA; 12 min incubation), Opal 620 (OP001004, Akoya Biosystems, Marlborough, Massachusetts, USA; 1/50, 8 min incubation); RICTOR (2114S, Cell Signaling Technology, Danvers, Massachusetts, USA; 1/25, 2 h incubation), UltraMap-antiRb HRP (760-4315, Roche Tissue Diagnostics, Oro Valley, Arizona, USA; 32 min incubation), Opal 690 (OP001006, Akoya Biosystems, Marlborough, Massachusetts, USA; 1/100, 8 min incubation); Ki67 (30–9) (790-4286, Roche Tissue Diagnostics, Oro Valley, Arizona, USA; ready-to-use solution, 20 min incubation), OmniMap-antiRb HRP (760-4311, Roche Tissue Diagnostics, Oro Valley, Arizona, USA; 12 min incubation), Opal 570 (OP001003, Akoya Biosystems, Marlborough, Massachusetts, USA; 1/200, 8 min incubation); pan-CKAE1-3 (NCL-L-AE1-3, Leica Biosystems, Wetzlar, Germany; 1/250, 28 min incubation), OmniMap-antiMs HRP (760-4310, Roche Tissue Diagnostics, Oro Valley, Arizona, USA; 12 min incubation), TSA-DIG (OP001007, Akoya Biosystems, Marlborough, Massachusetts, USA; 1/100, 20 min incubation) and Opal 780 (OP001008, Akoya Biosystems, Marlborough, Massachusetts, USA; 1/10, 1 h incubation). Following each detection with opal throughout the multiplex sequence, a denature step was included to denature any previous antibody binding using a heat-and pH-mediated approach (100℃ at pH 6). 3 drops of a DAPI counterstain (760-4196, Roche Tissue Diagnostics, Oro Valley, Arizona, USA) was applied for 16 min to each sample for nuclear detection.

Multispectral whole slide images were scanned on the PhenoImager imaging system (v1.0, Akoya Biosystems, Marlborough, Massachusetts, USA) using a 20x objective using Motif mode (acquisition version 1.0). The autofluorescence exposure time (ms) of an unstained lung control slide that had been treated on the Ventana Discovery Ultra with de-dewax and antigen retrieval steps only, was used to teach Inform (v6.0, Akoya Biosystems, Marlborough, Massachusetts, USA) the wavelength of autofluorescence in the lung for spectral unmixing. Core images were mapped, de-arrayed on Phenochart (v1.1.0, Akoya Biosystems, Marlborough, Massachusetts, USA) and stitched back together using Qupath (v0.4.3, University of Edinburgh, Edinburgh, UK) and the resulting component TMA images were added into Visiopharm (v2023.01.3.14018, Visiopharm, Hørsholm, Denmark) for image analysis. Core images were de-arrayed in Visiopharm using the TissueArray module to reveal patient IDs. For tissue segmentation, a bespoke deep learning algorithm was trained using the deep learning module using DAPI, CK, and autofluorescence channels to segment each core into tumor, stroma, and necrosis regions of interest (ROIs). For cell detection, another deep learning algorithm provided by Visiopharm was used as a template and was further trained using the deep learning module using the DAPI only channel to detect nuclei, nuclear boundaries, and background within ROIs, before post processing steps occurred to tidy up cellular labels. 12 TMA blocks of lung adenocarcinoma from the LATTICeA cohort as previously described (58), were analyzed, consisting of 1098 TMA cores covering 463 patients. Areas of each ROI, mean pixel intensities for each marker, and X Y coordinates were collected and exported for each core image for subsequent statistical analysis.

Raw single-cell data were imported from Visiopharm tab-separated value files. For each tissue core, cells were filtered to include only those located within "Tumor" or "Stroma" regions of interest as defined by the Visiopharm analysis. Intensity values for all markers were robustly normalized at the TMA level using the median and interquartile range (IQR) to mitigate the effect of outliers.

Normalized single-cell data were aggregated to the core level by calculating the median intensity for each marker within the "Tumor" and "Stroma" regions separately, as well as for the "Whole Core" (combining both regions). Case-level data were derived by averaging the core-level median intensities across all replicate cores for a given patient case.

## Polysome profiling

Transfected cells were scrapped and collected in a PBS solution supplemented with 100µg/mL of cycloheximide. Next, cells were lysed in hypotonic lysis buffer (2.5mM MgCl2, 5mM Tris-HCl (pH 7.5), 1.5mM KCl, 1x Complete™ protease inhibitor cocktail (Cat# 04693116001, Roche, Basel, Switzerland), 100µg/mL CHX, 0.5mM DTT), followed by sequential addition of RNasin® Ribonuclease Inhibitor (50 units) (N2515, Promega, Madison, Wisconsin, USA), Triton X-100 (0.5%) and Sodium Deoxycholate (0.5%). Samples were centrifuged to collect the cytosolic lysates, which were then loaded onto a non-linear sucrose gradient (5%-34%-55%) for ultracentrifugation (200,000 × g for 2 h at 4°C). Approximately 10% of the cytosolic lysate was reserved for total RNA extraction (input). Gradients were fractionated with the Piston Gradient Fractionator (BioComp Instruments Ltd., Fredericton, New Brunswick, Canada) to separate monosomes from polysomes while RNA content was monitored by UV absorbance (254nm). Finally, TriPure Isolation Reagent (Cat.# 11667165001, Roche, Basel, Switzerland) was added to the collected fractions and samples were snap-frozen and stored at -80°C for further extraction. RNA sequencing was performed on total RNA (input) and polysomal fractions (efficiently translated mRNA; associated with >3 ribosomes). Polysome profiling data were analyzed using a custom in-house R Shiny application. Translation efficiency (TE) was calculated as the ratio of the polysome peak area divided by the 40S peak area.

## TriPure extraction

RNA was extracted with TriPure Isolation reagent from Roche (Cat.# 11667165001, Roche, Basel, Switzerland) according to manufacturer’s instructions. Briefly, after TriPure and chloroform were added, samples were centrifuged and upper phase was collected. RNA was then precipitated with isopropanol and washed with 75% ethanol. After RNA was dried, it was dosed with NanoDrop 1000 Spectrophotometer (Thermo Fisher Scientific, Waltham, Massachusetts, USA).

## Transcriptomic analysis

Total RNA was extracted by TriPure extraction or by using the E.Z.N.A. Total RNA Kit I, as previously described. For RNA sequencing, RNA concentration was determined using the Quant-it™ RiboGreen Reagent and RNA Assay Kit (R11490, Thermo Fisher Scientific, Waltham, Massachusetts, USA). RNA quality and integrity was verified using the 5200 Fragment analyzer (Agilent Technologies, Santa Clara, California, USA). Total RNA was subjected to library preparation using the Illumina Stranded Total RNA Prep with Ribo-Zero Plus Kit (Cat# 20040525, Illumina, San Diego, California, USA). Libraries were then sequenced using an Illumina NovaSeq 6000 platform and the NovaSeq 6000 S4 Reagent Kit v1.5 (Cat# 20028312, Illumina, San Diego, California, USA). Sequencing data (reads) were aligned to a reference genome (Human: GRCh38; Mouse: GRCm39), quantified and normalized for further analysis between conditions.

## Proteomic analysis

Extracted cells were lysed with the PAC lysis buffer (5% SDS, 100mM Tris pH 8.5, 1mg/mL chloroacetamide, 1.5mg/mL Tris (2-carboxyethyl) phosphine). They were heated at 95°C for 30 min and then sonicated. Lysate was then added to a Kingfisher 96 well deep well plate and prepared for an 8 h digest protocol on the Kingfisher Duo (Thermo Fisher Scientific, Waltham, MA, USA). Lysate was added to Row G with MagReSyn HILIC beads and ACN is added to 70% final concentration. Rows D, E, and F are filled with 95% ACN and Rows B and C are filled with 70% EtOH. The digest buffer (1ug/mL MS grade trypsin in 50mM Triethyammonium bicarbonate) is added to Row A. The peptides were desalted using a C18 double disc in a P200 tip. The C18 filter was activated using MeOH and 0.1% TFA. The peptides were eluted using 50% ACN and 0.05% TFA after which they were dried and reconstituted in 0.1% TFA. The peptide concentration was measured using a Nanodrop to adjust for an equal input. Desalted peptides were then loaded onto 25cm Aurora Columns (IonOpticks, Fitzroy, Victoria, Australia ) using a RSLC nano uHPLC systems connected to a Fusion Lumos mass spectrometer (Thermo Fisher Scientific, Waltham, Massachusetts, USA). Peptides were separated by a 70 min linear gradient from 5% to 30% acetonitrile, 0.5% acetic acid. The mass spectrometer was operated in DIA mode, acquiring a MS 350-1650 Da at 120k resolution followed by MS/MS on 45 windows with 0.5 Da overlap (200-2000 Da) at 30k with a NCE setting of 28. The raw files were searched using the latest version of DIA-NN (v1.8 to v2.3) with the murine or human UniProt FASTA protein database. The precursor m/z range was set between 350-1650 and the fragment ion m/z range was set between 200-2000.

## Metabolomic analysis

2 × 10^5^ cells were seeded in 6-well plates and allowed to grow for 48 h. Cells were then washed twice with ice-cold PBS and polar metabolites were extracted in a MetOH-Acetonitrile-H_2_O (50-30-20) buffer for 5 min under gentle agitation. Collected supernatant was vigorously mixed for 10 min and centrifuged at 16,100 × *g* for another 10 min. All the steps were performed at 4 °C. Finally, supernatant was transferred to glass HPLC vials (Thermo Fisher Scientific, Waltham, Massachusetts, USA) and analysed using HPLC-MS.

Samples were analysed on a Q Exactive Orbitrap mass spectrometer (Thermo Fisher Scientific, Waltham, Massachusetts, USA) coupled with a Thermo Ultimate 3000 HPLC system. The HPLC setup consisted of a ZIC-pHILIC column (SeQuant, 150 × 2.1 mm, 5 µm, Merck KGaA, Darmstadt, Germany), with a ZIC-pHILIC guard column (SeQuant, 20 × 2.1 mm) and an initial mobile phase of 20% 20mM ammonium carbonate, pH 9.2, and 80% acetonitrile. Cell extracts (5 µL) were injected, and metabolites were separated over a 15 min mobile phase gradient, decreasing the acetonitrile content to 20%, at a flow rate of 200 μl/min and a column temperature of 45 °C. The total analysis time was 25 min. All metabolites were detected across a mass range of 75–1000 *m*/*z* using the Q Exactive mass spectrometer at a resolution of 35.000 (at 200 *m/z*), with electrospray ionisation (ESI) and polarity switching to enable both positive and negative ions to be determined in the same run. Lock masses were used, and the mass accuracy obtained for all metabolites was below 5 ppm. Data were acquired with Thermo Xcalibur software (Thermo Fisher Scientific, Waltham, Massachusetts, USA).

The peak areas of different metabolites were determined using Thermo TraceFinder 4.0 (Thermo Fisher Scientific, Waltham, Massachusetts, USA) software, where metabolites were identified by the exact mass of the singly charged ion and by known retention time on the HPLC column. Commercial standards of all metabolites detected had been analysed previously on this LC-MS system with the pHILIC column. Intracellular metabolites were normalised to protein content of the cells, measured at the end of the experiment by the BCA assay (Thermo Fisher Scientific, Waltham, Massachusetts, USA).

## LC-MS analysis of sphingosine and sphinganine

Analyses were conducted using a Thermo Fisher Scientific Ultimate 3000 binary UPLC system coupled to a Q Exactive Orbitrap mass spectrometer with a Heated Electrospray Ionization (HESI-II) source (Thermo Fisher Scientific, Waltham, Massachusetts, USA). Instrument control was performed with Xcalibur v4.3. Data were acquired in positive ion mode at a resolution of 70.000 at m/z 200, employing selected ion monitoring to improve detection sensitivity. Sphingosine (m/z 300.29) and Sphinganine (m/z 302.31) were monitored with a 0.7 m/z isolation offset. The automatic gain control target and maximum injection time were set to 2 × 10⁵ and 250 ms, respectively. Peaks were confirmed by matching expected accurate mass and fragmentation spectra. The retention time of Sphinganine was confirmed by reference to an authentic standard.

Chromatographic separation was achieved by gradient elution on a Waters ACQUITY Premier UPLC HSS T3 analytical column (150 × 2.1 mm, 1.8 µm) maintained at 45°C. The mobile phases consisted of water with 0.1% formic acid (A) and methanol with 0.1% formic acid (B) at a flow rate of 300 µL/min. The gradient started at 20% B, increased linearly to 95% B over 8 min, held for 1 min, then returned to initial conditions and equilibrated for 3 min, giving a total run time of 12 min. The injection volume was 5 µl. Raw data were processed in TraceFinder software (v4.1, Thermo Fisher Scientific, Waltham, Massachusetts, USA) to generate extracted ion chromatograms (XICs) using a mass tolerance of ±5 ppm. Relative quantification between sample groups was based on raw peak areas.

## Lipidomic analysis

Cell pellets were extracted in 200 µL of isopropanol, sonicated on ice-cold bath for 10 min, shaken for additional 10 min at 4°C, and finally centrifuged to precipitate proteins. Supernatants containing lipid extracts were then individually transferred to new vials and separated on a Dionex UltiMate 3000 LC System (Thermo Fisher Scientific, Waltham, Massachusetts, USA) using a Kinetex C18 EVO 2.6µM, 100A, 150 x 0.3 mm LC Column following the sequence:

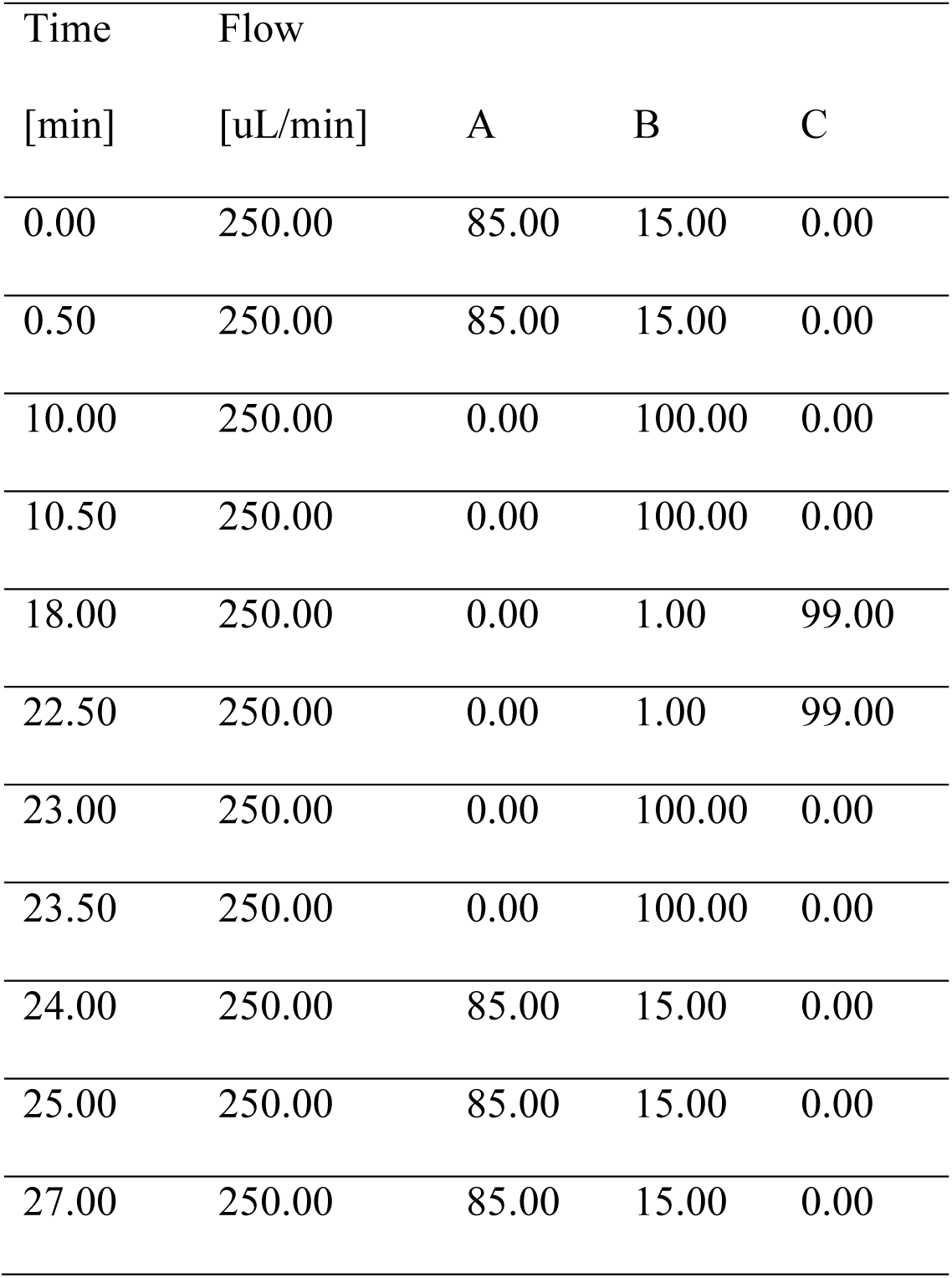

| Time | Flow |  |  |  |
| --- | --- | --- | --- | --- |
| [min] | [μL/min] | A | B | C |
| 0.00 | 250.00 | 85.00 | 15.00 | 0.00 |
| 0.50 | 250.00 | 85.00 | 15.00 | 0.00 |
| 10.00 | 250.00 | 0.00 | 100.00 | 0.00 |
| 10.50 | 250.00 | 0.00 | 100.00 | 0.00 |
| 18.00 | 250.00 | 0.00 | 1.00 | 99.00 |
| 22.50 | 250.00 | 0.00 | 1.00 | 99.00 |
| 23.00 | 250.00 | 0.00 | 100.00 | 0.00 |
| 23.50 | 250.00 | 0.00 | 100.00 | 0.00 |
| 24.00 | 250.00 | 85.00 | 15.00 | 0.00 |
| 25.00 | 250.00 | 85.00 | 15.00 | 0.00 |
| 27.00 | 250.00 | 85.00 | 15.00 | 0.00 |

Mobile phases consist of (A) H_2_O - 5mM ammonium formate, 0.1% Formic acid, (B) 60% Acetonitrile: 40% Methanol - 5mM ammonium formate, 0.1% Formic acid, and (C) Isopropanol - 5mM ammonium formate, 0.1% Formic acid.

Mass spectrometry data were acquired on a Q Exactive™ Plus Hybrid Quadrupole-Orbitrap™ Mass Spectrometer (Thermo Fisher Scientific, Waltham, Massachusetts, USA) using data dependent acquisition in positive and negative ion mode. The parameters used were: **Full MS:** Resolution: 70,000; AGC target: 1e6; Max IT: 100 ms; Scan range: 90 to 1350 m/z. **dd-MS^2:^** Resolution: 17,500; AGC target: 1e5; Max IT: 120 ms; Loop count: 5; TopN: 5; Isolation window: 1.5 m/z.

Data were processed using compound discovery (Thermo Fisher Scientific, Waltham, Massachusetts, USA) and lipids identification was performed using Lipidex (59).

## MALDI-MSI Spatial Lipidomics

Fresh-frozen tissue samples were sectioned at a thickness of 10µm using a Microm HM525 NX cryostat (Thermo Fisher Scientific, Waltham, Massachusetts, USA) maintained at −20°C. Sections were thaw-mounted onto conductive IntelliSlides (Bruker, Billerica, Massachusetts, USA), dried for 10 min at room temperature in a vacuum desiccator, vacuum-sealed, and stored at −80°C until analysis. For MALDI mass spectrometry imaging in negative ion mode, a matrix solution of 2,5-dihydroxyacetophenone (DHAP; 5mg/mL) prepared in a 2:1 (v/v) mixture of chloroform and methanol was applied using an HTX M3+ sprayer (HTX Technologies, Chapel Hill, North Carolina, USA). Spraying was performed at a flow rate of 0.125mL/min, with the nozzle temperature set to 30°C, a nozzle velocity of 1,350 mm/min, and a total of 10 spray passes. For positive ion mode analysis, norharmane (7mg/mL) dissolved in 2:1 chloroform:methanol (v/v) was deposited under identical conditions, except that 12 spray passes were applied. MALDI–MSI experiments were conducted using a timsTOF fleX MALDI mass spectrometer (Bruker, Billerica, Massachusetts, USA). Data were acquired in positive ion mode over an m/z range of 300–1800 and in negative ion mode over an m/z range of 250–2600. Spatial resolutions 20 × 20µm were used, with 200 laser shots per pixel and a laser repetition rate of 10kHz. Image acquisition was performed using FlexImaging (v7.0, Bruker, Billerica, Massachusetts, USA). MSI datasets were processed in SCiLS Lab (v2026, Bruker, Billerica, Massachusetts, USA) using a mass tolerance of ± 5ppm and with no denoising applied. Regions of interest and spectral data were exported to MetaboScape (v2025, Bruker, Billerica, Massachusetts, USA), where metabolite annotation was performed using a built-in custom lipid database. Following MSI acquisition, matrix residues were removed by immersing the slides in 100% methanol. Tissue morphology was subsequently assessed by H&E staining, allowing to define *Healthy* and *Tumor* subregions of interest within each tissue section.

For downstream analysis of positive- and negative-ionization MALDI-MSI spatial lipidomics data, spatially-resolved total ion current (TIC)-normalized abundance data for all lipid species of interest detected for each polarity were first exported as CSV files within SCiLS Lab 2025 (Bruker, Billerica, Massachusetts, USA), and further analyzed within the *R* framework (www.R-project.org). TIC-normalized data for each lipid species across each tissue section (N = 3 for both *KP* and *KPR^OE^*^)^ were first aggregated within each of the previously defined subregions of interest (n = 9-13 *Healthy* subregions and n = 19-42 *Tumor* subregions per section), and annotated accordingly to either region type. Aggregation was performed simply by averaging the TIC-normalized data across all pixels in each of these subregions. The subregion-aggregated counts for each lipid species were further log2-transformed, with any zeros first set to 50% of the lowest non-zero value for the corresponding lipid species across all subregions and tissue sections (i.e. regardless of genotype or region type), to avoid numerical overflow errors upon log2-transformation. Statistical analysis was then performed based on a linear categorical model applied on a per-lipid-species basis to these log2-transformed, subregion-aggregated counts. This was done with *R*’s function *lm()*, using a model of the form:

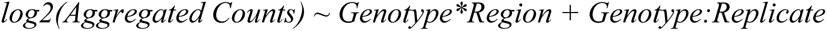

with (a) each lipid species’ log2-transformed, subregion-aggregated counts representing the response variable, (b) both the genotype (*KP/KPR^OE^*^)^ and region type (*Healthy/Tumor*) treated as interacting categorical predictors, and (c) the tissue section of origin treated as a blocking factor (via the rightmost term, where *Replicate* = *A/B/C*, corresponding to the N = 3 tissue sections available for each genotype). Based on the above model, the interaction coefficient *GenotypeKPR^OE:^RegionTumor* directly represents, for each lipid species, an estimate of the difference *Δlog_2_FC* in *Tumor vs. Healthy* log2-fold-changes between the *KPR^OE^* and *KP* conditions, with these estimates further accounting (via the blocking term) for the paired nature of the *Tumor vs. Healthy* data for each tissue section (namely, the fact that different tissue sections may present different *Healthy* baseline levels for each lipid species). The differences *Δlog_2_FC* can therefore be interpreted as baseline-corrected fold-changes in lipid-species levels between *KPR^OE^* and *KP* tumors, with positive/negative *Δlog2FC* values thus indicating higher/lower baseline-corrected levels for a given lipid species in *KPR^OE^* relative to *KP*, respectively. Effect sizes (indicating the magnitude and direction of the baseline-corrected fold-changes *Δlog_2_FC*) and p-values (indicating the significance levels for these baseline-corrected fold-changes) were extracted for each lipid species based on the above interaction coefficient, with p-values further being subject to false discovery rate (FDR) adjustment based on the Benjamini-Hochberg method, considering all lipid species included in the analysis for each polarity. A lipid species was considered to be significantly altered between *KPR^OE^* and *KP* tumors if its FDR-adjusted p-value was below 0.05, and significantly-altered lipid species were further ranked based on the absolute magnitude of the product *-log_10_(p-value)* x *Δlog_2_FC*, separately for each enrichment direction (i.e. for either positive or negative values of *Δlog_2_FC*). For dot-plot presentation, average levels of the log2-transformed, subregion-aggregated counts for each significantly-altered lipid species were first determined across all *Healthy* subregions in each tissue section. The latter were then subtracted from the log2-transformed subregion-aggregated data for that lipid species in the corresponding tissue section, so as to display baseline-corrected, log2-transformed, subregion-aggregated counts. For heatmap presentation, the above baseline-corrected, log2-transformed, subregion-aggregated counts for each lipid species were averaged across all *Tumor* subregions in each tissue section, and these average baseline-corrected, log2-transformed *Tumor* levels were then transformed into Z-score distributions for each lipid species across all tissue sections (to enable comparison across lipid species, with the latter further averaged across the 3 replicates for each condition (*KP* or *KPR^OE^*^)^).

## Bioinformatic analysis

Bioinformatic analysis were performed on in-house and publicly available transcriptomic and proteomic datasets. All our generated datasets were quality controlled, normalized and analyzed using standard pipelines. Differential expression was conducted using Limma (v3.60.4) on counts normalized to log10(counts per million) using edgeR (v4.2.1) https://doi.org/10.1093/nar/gkaf018 for library size normalization. For patient analysis, HTSeq counts for the GDC TCGA LUAD cohort were downloaded from Xena Browser (https://xenabrowser.net) and subsequently normalized to log10(counts per million) using edgeR (v4.2.1) https://doi.org/10.1093/nar/gkaf018 for library size normalization. A score for HIF1B signature (n = 100) was obtained using the GSVA R package (v1.48.0) https://doi.org/10.1186/1471-2105-14-7 and enrichment was assessed using the ssGSEA method with a Poisson kernel (60). Samples were subsequently stratified into groups according to their enrichment score for the HIF1B signature. Samples with a score superior to the median of the distribution were assigned to the “HIF1B^high”^ group and samples with a score inferior to the median of the distribution were assigned to the “HIF1B^low”^ group. The subsequent gene set enrichment analyses were performed using the Broad Institute GSEA software (v4.2.2) and the gene sets from MsigDB (v2022).

## Statistical analysis

GraphPad PRISM software (v8.4.3, GraphPad Software Inc, San Diego, California, USA) was used for statistical analysis.

## Data availability

Sequencing data have been deposited on the Gene Expression Omnibus database. For proteomics, the raw files and the MaxQuant search results files have been deposited as partial submission to the ProteomeXchange Consortium via the PRIDE partner repository (61). For lipidomics, the raw LC-MS/MS files have been deposited on the Metabolomics Workbench repository (62).

## Acknowledgments

Authors thank Maxence Casamento, Gwendoline Faye, Fiona Ballantyne and Catherine Ficken for providing technical assistance. We are grateful to the GIGA-animal, immunohistology, proteomics, imaging, genomics, bioinformatics and viral vector facilities, for their assistance. We thank the VIB Transgenic Core Facility for technical assistance with generation of *R26-mRictor^tg^* mouse line.

This study was supported by research grants from the FNRS (MIS F.4005.24 and CDR J.0018.25 awarded to A. Blomme, EOS O.0020.22 awarded to P. Close), the Belgian foundation against Cancer (2024-146 awarded to A. Blomme, 2024-187 awarded to D. Cataldo and 2020-068, 2024-148 awarded to P. Close), the Walloon Excellence in Life Sciences and Biotechnology (WELBIO CR-2022 A-03 awarded to P. Close), the European Union’s Horizon 2020 research and innovation program (Marie Sklodowska-Curie grant agreement No 101029147 awarded to A. Blomme) and the University of Liege. We are also grateful to the “Foundation Leon Fredericq”, the “KU Leuven Opening the Future campaign” and the “King Baudouin Foundation” for their financial support. S.-M. Fendt acknowledges funding from FWO Projects, Fonds Baillet Latour, Francqui Stichting, Foundation ARC, KU Leuven Methusalem (METH/26/009), Stichting tegen Kanker, Wereld Kanker Onderzoek Fonds (WKOF), as part of the World Cancer Research Fund International grant program (IIG_FULL_2023_001) and the Interuniversity BOF (iBOF) program. J. Le Quesne acknowledges the support of the NHS Greater Glasgow and Clyde Biorepository, the Glasgow Tissue Research Facility, and the CRUK Scotland Institute Histology Facility. M. Paque, N. An, and D. Vanneste are FNRS Research Fellow. A. Aliju is supported by a Televie fellowship. A. Blomme and M. Herfs are FNRS Research Associate. P. Close is FNRS Research Director.

## Author contributions

M.P., P.C. and A.B. designed the study and analyzed experimental data; M.P., N.A., M.L., J. F-G., I.P., P.C. and A.B. performed software and bioinformatic analyses; M.P., N.A., A.A., E.R., D.V., D.H., J.M.J., C.P., T.M., F.P., R.V., C.S., K.E., P.R., E.S., R.L.B., L. O-J., S.M., M.Pl., A.B. performed experiments; D.S., T.M., D.C., E.dV., S.G., G.B., J.L.Q., M.H., J.M., R.B., J.V.S., S-M.F., A.vK., P.C. and A.B. provided resources; M.P., P.C. and A.B. wrote the paper. All authors discussed the results and commented on the manuscript.

Correspondence and requests for materials should be addressed to;.

---

**Supplementary Figure 1.**
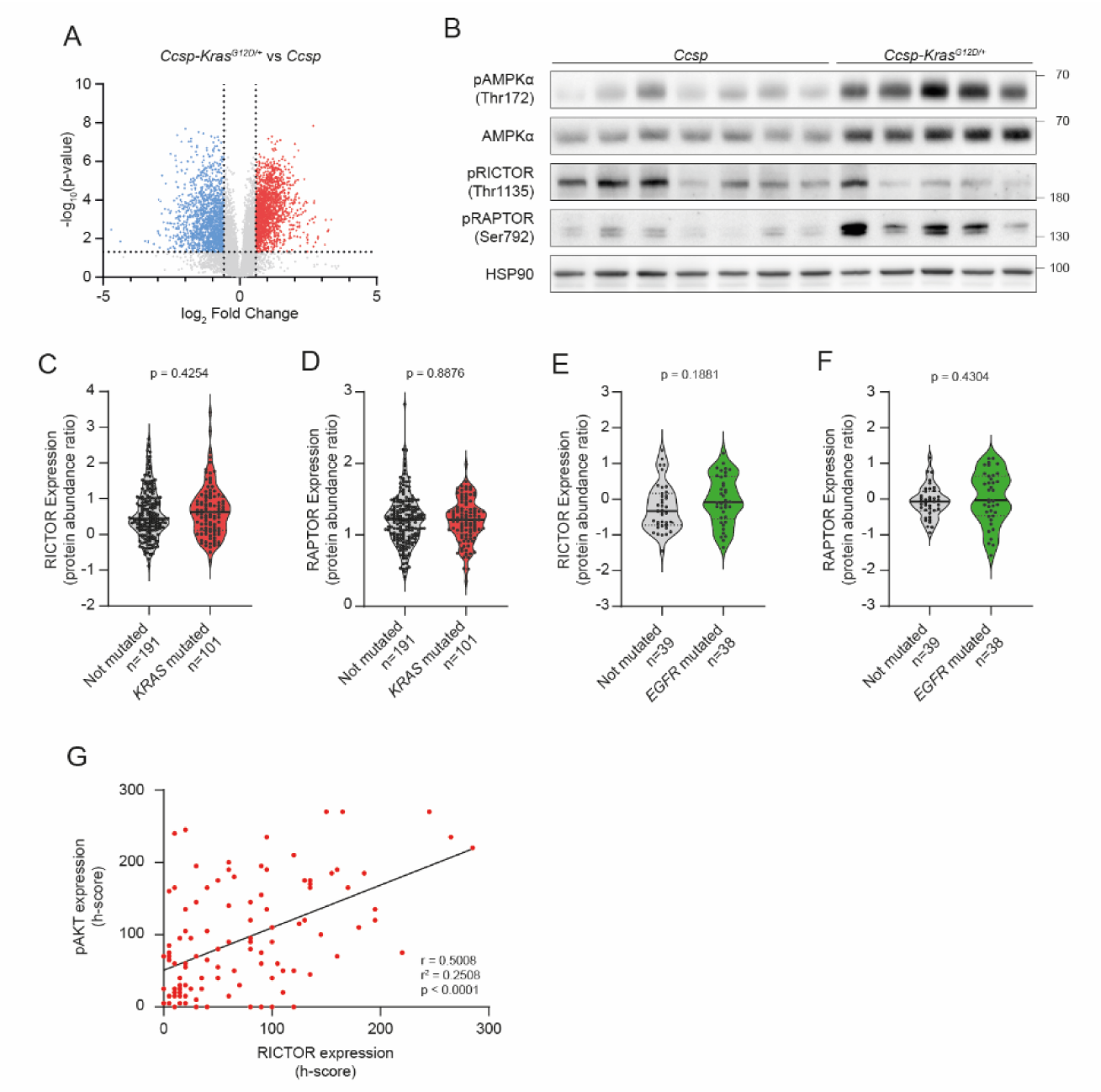
A. Volcano plot displaying the differentially expressed proteins in *Ccsp-Kras^G12D/+^* (n = 5) lung tumors when compared to *Ccsp* (n = 4) lungs. Significantly increased or decreased proteins (p-value < 0.05) are highlighted in red and blue, respectively. **B.** Western blot (WB) analysis of phosphorylated-AMPKα Thr172 (pAMPKα Thr172), AMPKα, phosphorylated-RICTOR Thr1135 (pRICTOR Thr1135) and phosphorylated-RAPTOR Ser792 (pRAPTOR Ser792) protein expression in lungs isolated from *Ccsp* and *Ccsp-Kras^G12D/+^* mice. HSP90 was used as a sample loading control. Representative image from one experiment with 7 animals for *Ccsp* and 5 for *Ccsp-Kras^G12D/+^*^.^ **C, D.** Violin plots showing RICTOR (**C**) and RAPTOR (**D**) protein abundance in LUAD samples stratified by *KRAS* mutational status (TCGA dataset, n = 292, Mann-Whitney U test). **E, F.** Violin plots of RICTOR (**E**) and RAPTOR (**F**) protein abundance in LUAD samples stratified by *EGFR* mutational status (CPTAC dataset, n = 77, Mann-Whitney U test). **G.** Scatter plot showing the correlation between RICTOR and pAKT Ser473 protein levels, as quantified by h-score, in lung cancer biopsies (Cologne cohort, n = 107, Pearson correlation).

**Supplementary Figure 2.**
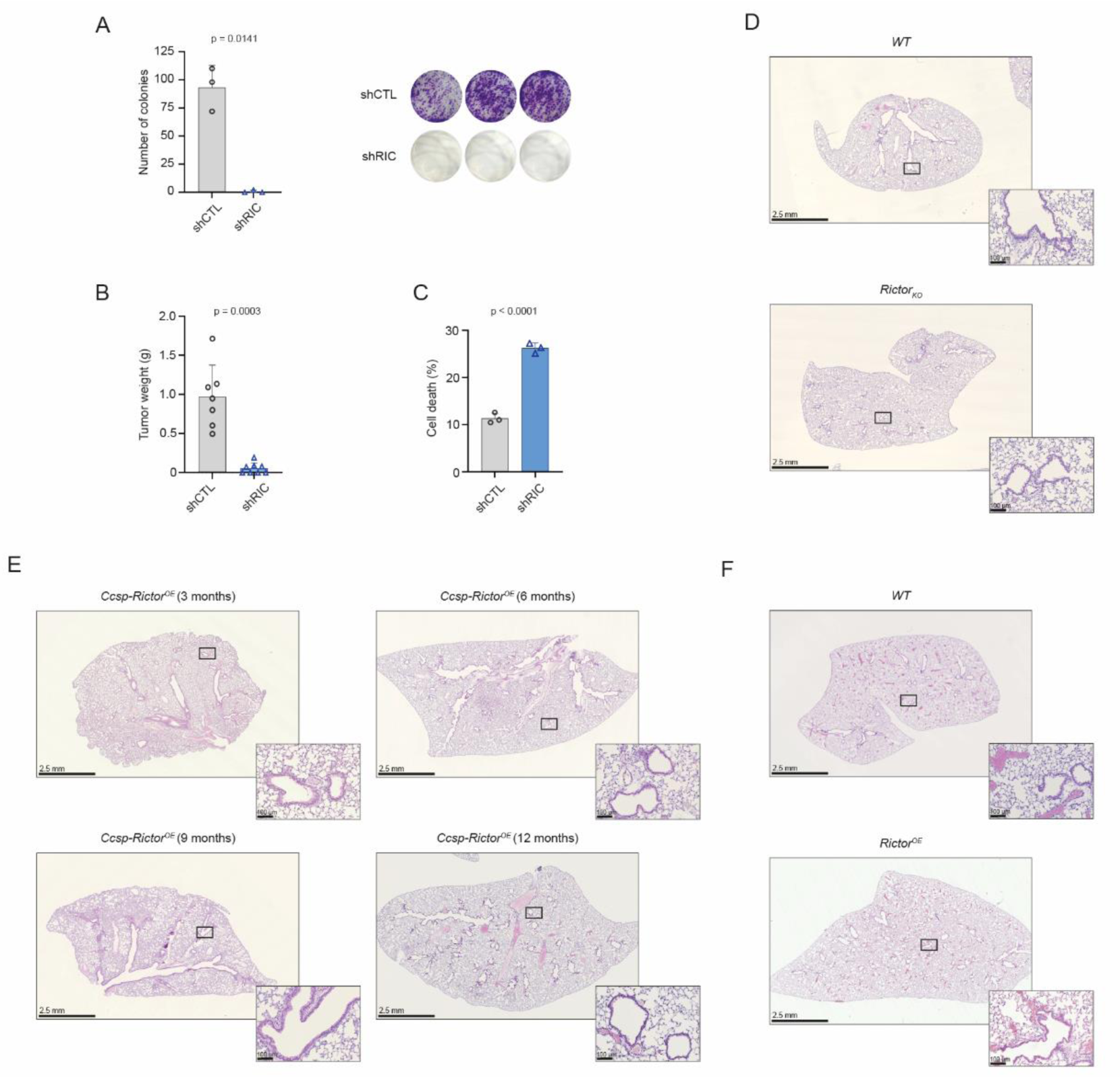
A. Quantification of colony formation assays performed with control (shCTL) and RICTOR-depleted (shRIC) A549 cells (mean ± SD, n = 3 technical triplicates from a representative biological experiment (performed in independent biological triplicates), Welch’s t-test). **B.** Endpoint tumor weights of shCTL and shRIC A549 xenografts grown in immunodeficient mice (mean ± SD, n = 7 tumors for shCTL and n = 8 tumors for shRIC, Mann-Whitney U test). **C.** Annexin V-PI staining (cell death) in shCTL and shRIC A549 cells (mean ± SD, n = 3 technical triplicates from a representative biological experiment (performed in independent biological triplicates), Welch’s t-test). **D.** Representative H&E-stained lung sections from WT and Rictor_KO_ mice. Scale bar represents 2.5 mm (left) and 100 µm (right). **E.** Representative H&E-stained lung sections from *Ccsp-Cre;Rictor^OE^* (Ccsp-Rictor^OE)^ mice at 3, 6, 9 and 12 months. Scale bar represents 2.5 mm (left) and 100 µm (right). **F.** Representative H&E-stained lung sections from WT and Rictor^OE^ mice. Scale bar represents 2.5 mm (left) and 100 µm (right).

**Supplementary Figure 3.**
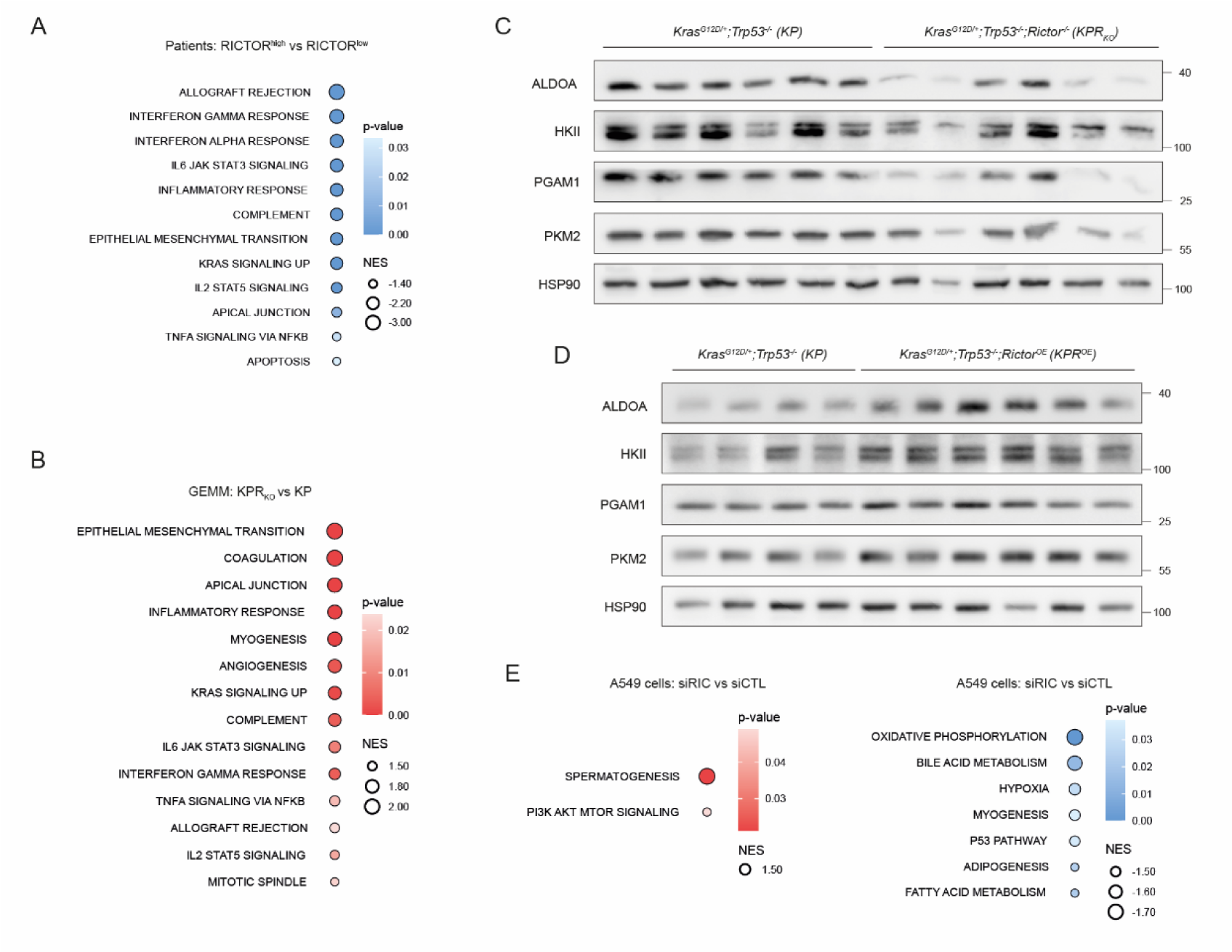
A. GSEA performed on proteomics of lung cancer samples from RICTOR high patients (RIC^high,^ n = 22, upper quartile) compared to RICTOR low patients (RIC^low,^ n = 22, lower quartile). Data was retrieved from Lehtiö et al (63). Shown are the significantly down-regulated Hallmark pathways ranked by NES and colored by significance (p-value). **B.** GSEA performed on proteomics of macro-dissected lung tumors isolated from KPR_KO_ mice (n = 5) compared to KP mice (n = 6). Shown are the significantly enriched Hallmark pathways ranked by NES and colored by significance (p-value). **C.** WB analysis of ALDOA, HKII, PGAM1 and PKM2 protein expression in macro-dissected lungs isolated from KP and KPR_KO_ mice. HSP90 was used as a sample loading control. Representative image from one experiment with 6 animals per group. **D.** WB analysis of ALDOA, HKII, PGAM1 and PKM2 protein expression in macro-dissected lungs isolated from KP and KPR^OE^ mice. HSP90 was used as a sample loading control. Representative image from one experiment with 4 animals for KP and 6 for KPR^OE.^ **E.** GSEA performed on proteomics of A549 cells silenced for RICTOR (siRIC) compared to control cells (siCTL). Shown are the significantly enriched (left) and downregulated (right) Hallmark pathways ranked by NES and colored by significance (p-value).

**Supplementary Figure 4.**
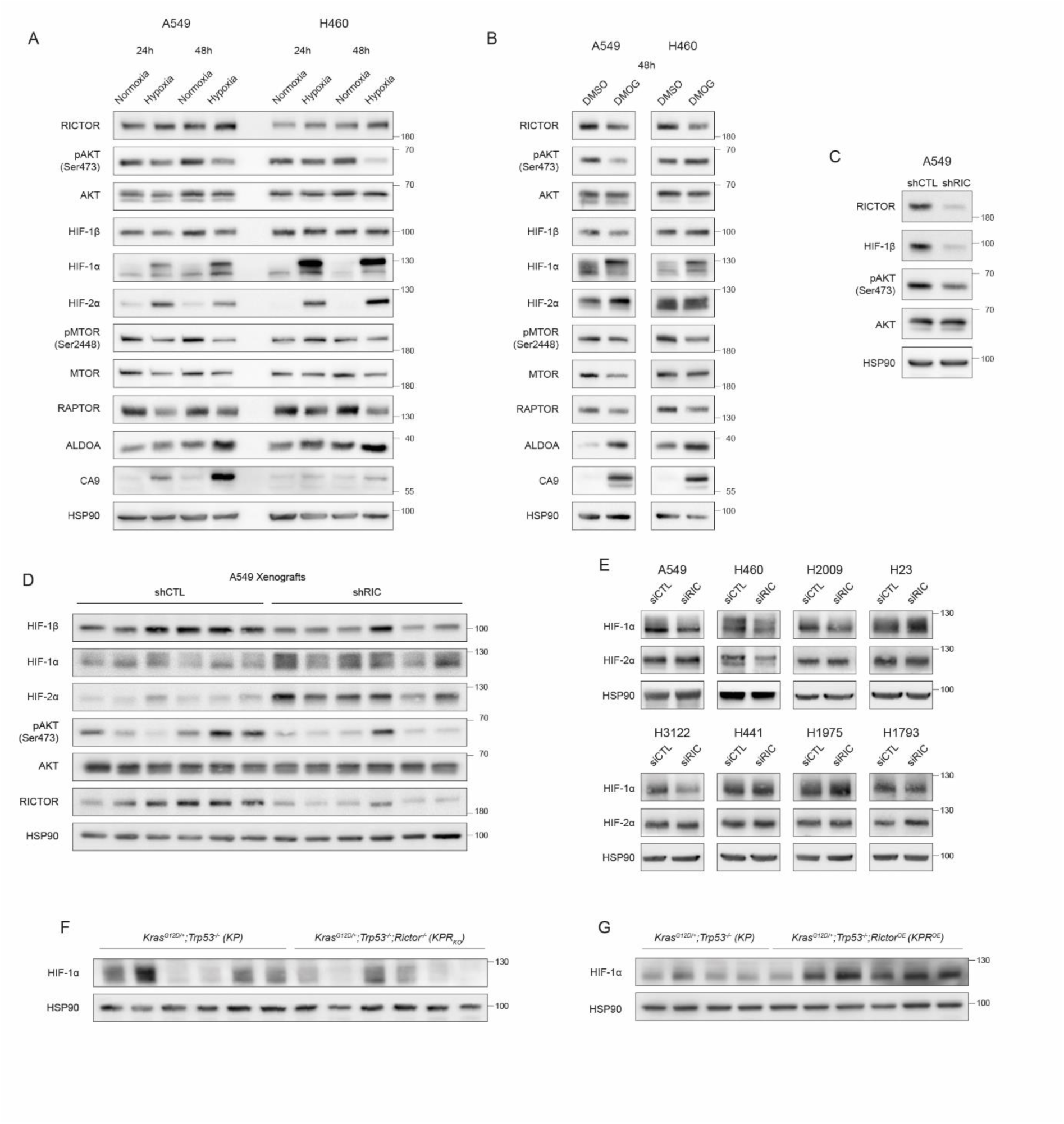
A. WB analysis of RICTOR, pAKT Ser473, AKT, HIF-1β, HIF-1α, HIF-2α, pMTOR Ser2448, MTOR, RAPTOR, ALDOA and CA9 protein expression in A549 and H460 cell lines grown under normoxia or hypoxia conditions for 24h and 48h. HSP90 was used as a sample loading control. Representative image from three independent biological experiments. **B.** WB analysis of RICTOR, pAKT Ser473, AKT, HIF-1β, HIF-1α, HIF-2α, pMTOR Ser2448, MTOR, RAPTOR, ALDOA and CA9 protein expression in A549 and H460 cell lines treated with DMOG (HIF-1α stabilizer, 1mM) for 48h. HSP90 was used as a sample loading control. Representative image from three independent biological experiments. **C.** WB analysis of RICTOR, HIF-1β, pAKT Ser473 and AKT protein expression in control (shCTL) and RICTOR-depleted A549 cells (shRIC). HSP90 was used as a sample loading control. Representative image from three independent biological experiments. **D.** WB analysis of HIF-1β, HIF-1α, HIF-2α, pAKT Ser473, AKT and RICTOR protein expression in shCTL and shRIC A549 xenografts grown in immunodeficient mice. HSP90 was used as a sample loading control. Representative image from one experiment with 6 animals per group. **E.** WB analysis of HIF-1α and HIF-2α protein expression following RICTOR silencing in multiple human lung cancer cell lines. HSP90 was used as a sample loading control. Representative image from three independent biological experiments. **F.** WB analysis of HIF-1α protein expression in macro-dissected lung tumors isolated from KP and KPR_KO_ mice. HSP90 was used as a sample loading control. Representative image from one experiment with 6 animals per group. **G.** WB analysis of HIF-1α protein expression in macro-dissected lung tumors isolated from KP and KPR^OE^ mice. HSP90 was used as a sample loading control. Representative image from one experiment with 4 animals for KP and 6 for KPR^OE.^

**Supplementary Figure 5.**
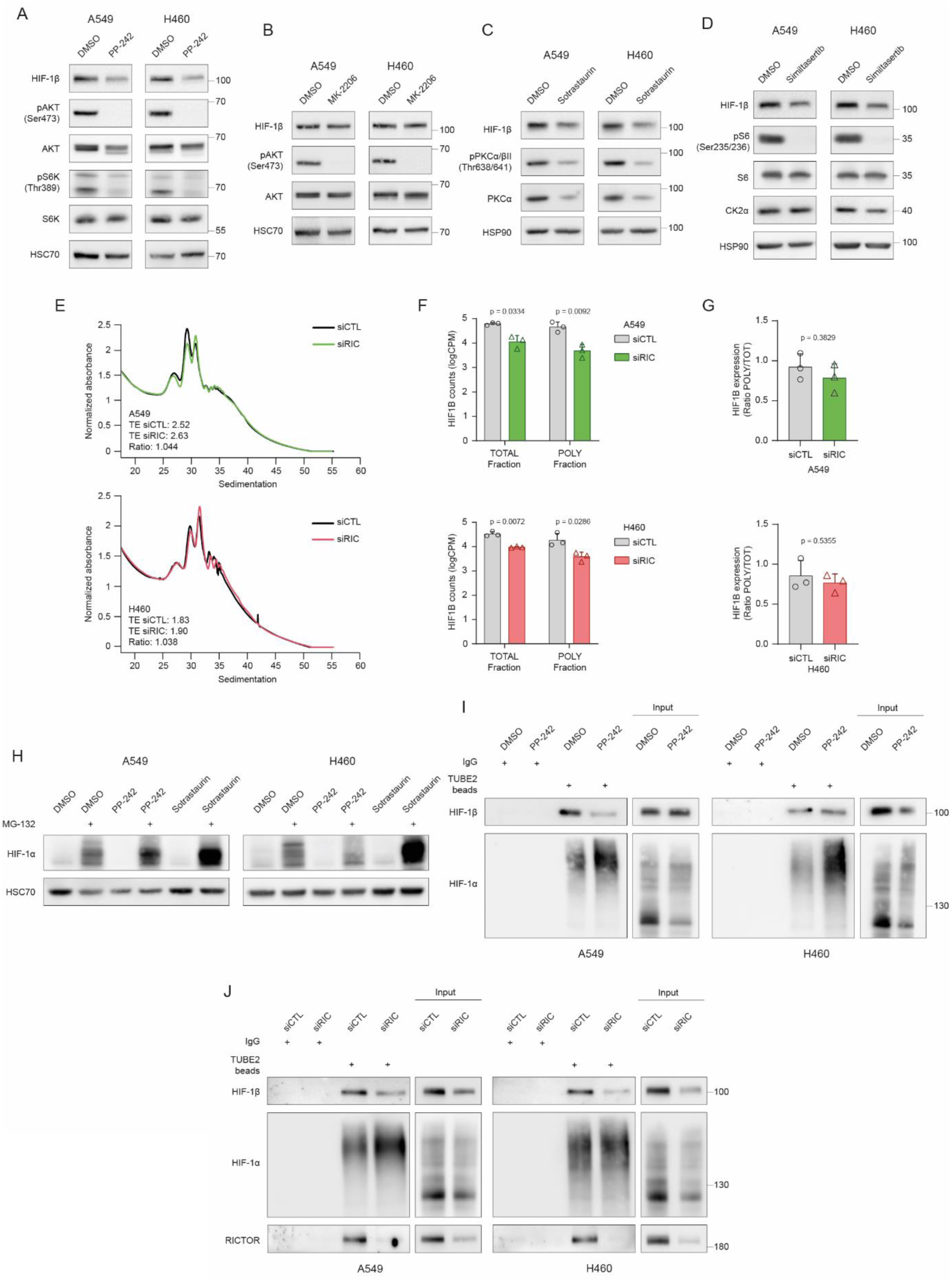
A. WB analysis of HIF-1β, pAKT Ser473, AKT, phosphorylated-P70S6K Thr389 (pS6K Thr389) and P70S6K (pS6K) protein expression following 8h treatment with PP-242 (10µM) or vehicle control in A549 and H460 cells. HSC70 was used as a sample loading control. Representative image from three independent biological experiments. **B.** WB analysis of HIF-1β, pAKT Ser473 and AKT protein expression following 8h treatment with MK-2206 (2µM) or vehicle control in A549 and H460 cells. HSC70 was used as a sample loading control. Representative image from three independent biological experiments. **C.** WB analysis of HIF-1β, phosphorylated-PKCα/β Thr638/641 (pPKCα/β Thr638/641) and PKCα protein expression following 8h treatment with sotrastaurin (50µM) or vehicle control in A549 and H460 cells. HSP90 was used as a sample loading control. Representative image from three independent biological experiments. **D.** WB analysis of HIF-1β, phosphorylated-S6 Ser235/236 (pS6 Ser235/236), S6 and CK2α protein expression following 8h treatment with silmitasertib (50µM) or vehicle control in A549 and H460 cells. HSP90 was used as a sample loading control. Representative image from three independent biological experiments. **E**. Polysome profiles of control (siCTL) and RICTOR-depleted (siRIC) A549 (above) and H460 (below) cells. Translational efficiency (TE) was calculated for each profile, and the TE ratio of RICTOR-silenced versus control cells is represented. Representative profile from three independent biological experiments for each cell line. **F.** *HIF1B* mRNA abundance (in CPM (Counts per Million)) determined by transcriptomic analysis in total and polysomal fractions from siCTL and siRIC A549 (above) and H460 (below) cell lines (n = 3 independent biological experiments, mean ± SD, Welch’s t-test). **G.** Ratio of polysomal to total *HIF1B* mRNA levels in siCTL and siRIC A549 (above) and H460 (below) cell lines (n = 3 independent biological experiments, mean ± SD, Welch’s t-test). **H.** WB analysis of HIF-1α protein expression in A549 and H460 cells treated with DMSO, PP-242 (10µM) or sotrastaurin (50µM), in the presence or absence of MG-132 (20µM). HSC70 was used as a sample loading control. Representative image from three independent biological experiments. **I.** WB analysis of HIF-1β and HIF-1α following immunoprecipitation of ubiquitinated proteins (TUBE2 beads) in A549 and H460 cells treated with DMSO or PP-242 (10µM) for 24h in presence of MG-132 (20µM). Representative image from a single biological experiment. **J.** WB analysis of HIF-1β, HIF-1α and RICTOR following immunoprecipitation of ubiquitinated proteins (TUBE2 beads) in siCTL and siRIC A549 and H460 cells, in presence of MG-132 (20µM). Representative image from three independent biological experiments.

**Supplementary Figure 6.**
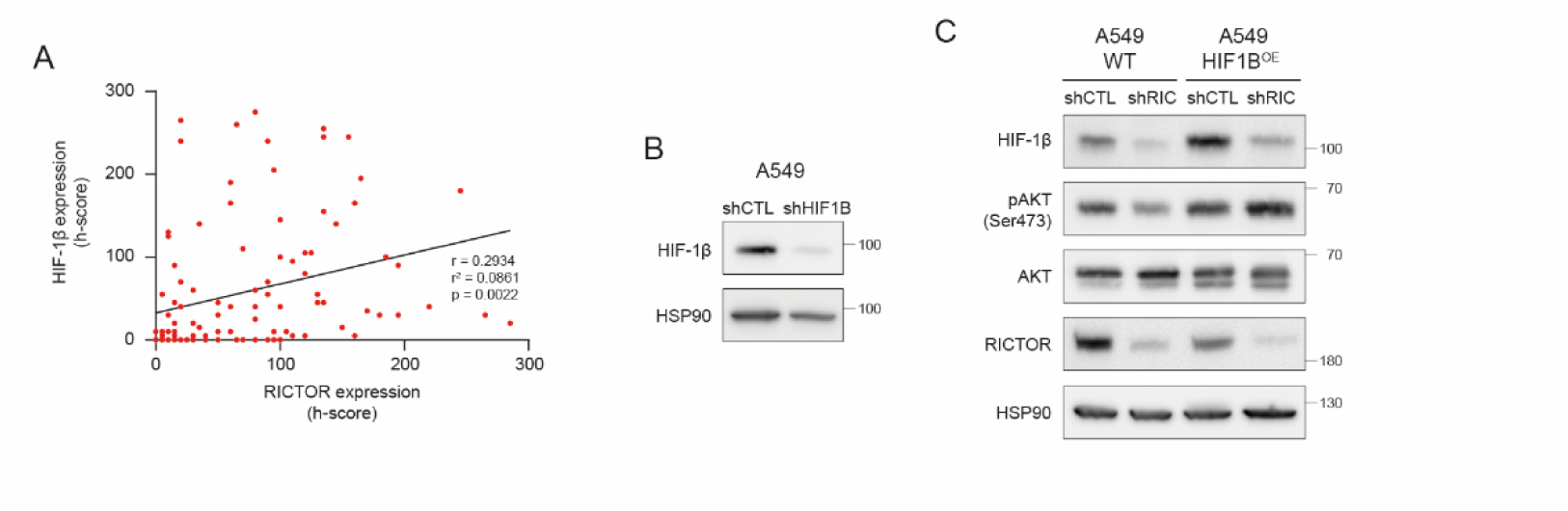
A. Scatter plot showing the correlation between RICTOR and HIF-1β protein levels (quantified by h-score) in lung cancer biopsies (Cologne cohort, n =107, Pearson correlation). **B.** WB analysis of HIF-1β protein expression in control or HIF1B-depleted (shHIF1B) A549 cells. HSP90 was used as a sample loading control. Representative image from three independent biological experiments. **C.** WB analysis of HIF-1β, pAKT Ser473, AKT and RICTOR protein expression in WT or HIF1B-overexpressing A549 cells (HIF1B^OE)^ depleted or not for RICTOR (shCTL and shRIC). HSP90 was used as a sample loading control. Representative image from three independent biological experiments.

**Supplementary Figure 7.**
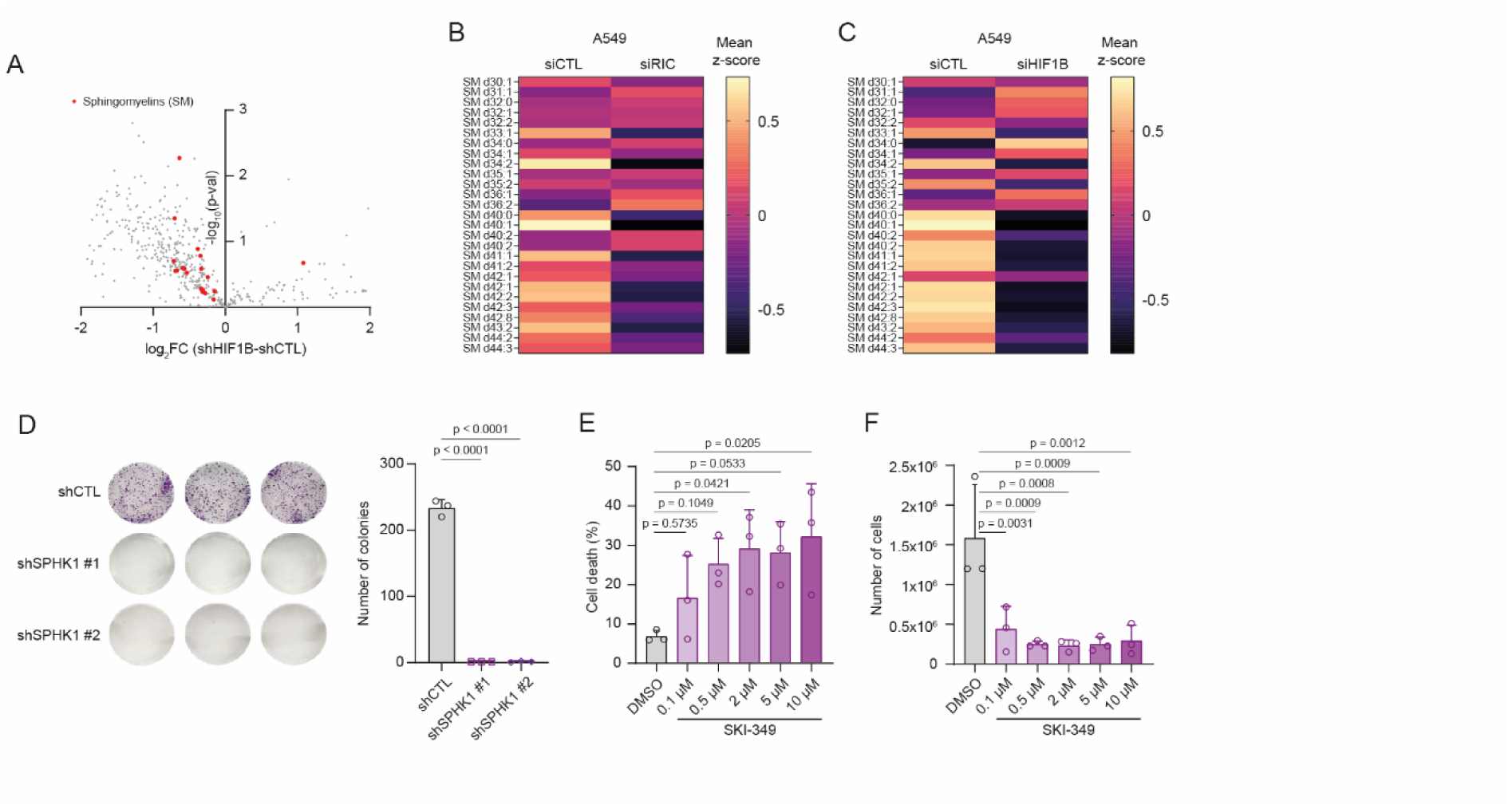
A. Volcano plot of the differentially abundant lipids in shHIF1B versus shCTL A549 cells. Sphingomyelins are highlighted in red (n = 3 independent biological experiments, Student t-test). **B, C.** Heatmaps illustrating the differential abundance of sphingomyelin species in A549 cells following RICTOR (siRIC) (**B**) or HIF1B (siHIF1B) (**C**) depletion (mean Z-scores, n = 3 independent biological experiments per group). **D.** Quantification of colony formation assays performed with control (shCTL) and SPHK1-depleted (shSPHK1) A549 cells (mean ± SD, n = 3 technical triplicates from a representative biological experiment (performed in independent biological triplicates), one-way ANOVA with Dunnet’s correction test). **E, F.** Annexin V-PI staining (cell death) (**E**) and cell number (**F**) in A549 cells treated with increasing concentrations of SKI-349 (mean ± SD, n = 3 independent biological experiments, one-way ANOVA with Dunnet’s correction test).

