## Supplementary Tables for "mTORC2 stabilizes HIF-1β to coordinate metabolic adaptation in lung cancer"

1) Suppl. Table 1: shRNA used in this study

| Name of the shRNA | Vector backbone | Stock concentration and concentration used | Reference | Antibiotic |
|---|---|---|---|---|
| **shRICTOR #5** | shRictor#5 | 539ng/µL – 12 µg | ML-21.0224 | Puromycin |
| **shHIF1B #1** | pLV-U6 shRNA ARNT #1 | 150ng/µL – 12µg | ML-22.0127 GVV_1799 | Puromycin |
| **shHIF1B #2** | pLV-U6 shRNA ARNT #2 | 300ng/µL – 12µg | ML-22.0128 GVV_1800 | Puromycin |
| **OE HIF1B** | pLV EF1A HA-hARNT WT DNA plasmid 18 E7 | 375ng/µL – 12µg | PL-23.0130 ML-23.0173 | Hygromycin |
| **shSPHK1 #1** | pLV U6 hSPHK1#1 | 494ng/µL – 12µg | PL-25.0216 ML-25.0292 | Puromycin |
| **shSPHK1 #11**[1] | pLV U6 hSPHK1#11 | 315ng/µL – 12µg | PL-25.0219 ML-25.0295 | Puromycin |

2) Suppl. Table 2: Primers used in this study

Primers were ordered on integrated DNA technologies (IDT) following verification of off-targeting on the NCBI primer (Blast) site.

| Primer name | Sequence | Company |
|---|---|---|
| **GAPDH** | Forward Sequence: GTCTCCTCTGACTTCAACAGCG<br>Reverse Sequence: ACCACCCTGTTGCTGTAGCCAA | Integrated DNA Technologies (Coralville, Iowa, USA) |
| **HIF1β** | Forward Sequence: CTGTCATCCTGAAGACCAGCAG<br>Reverse Sequence: CTGGTTCTCATCCAGAGCCATTC | Integrated DNA Technologies (Coralville, Iowa, USA) |

3) Suppl. Table 3: Antibodies used in this study

All antibodies are diluted 1/1000 in 5% BSA.

| | Target | Reference | Supplier + Catalogue number |
|---|---|---|---|
| **1.** | RICTOR | #2140 | Cell Signaling Technology (Danvers, Massachusetts, USA) |

[1] For representation matters, #11 has been replaced by #2 in figures

| | | | |
|---|---|---|---|
| **2.** | Phospho-RICTOR (Thr1135) | #3806 | Cell Signaling Technology |
| **3.** | RAPTOR | #48648 | Cell Signaling Technology |
| **4.** | Phospho-RAPTOR (Ser792) | #2083 | Cell Signaling Technology |
| **5.** | mTOR | #2983 | Cell Signaling Technology |
| **6.** | Phospho-mTOR (Ser2448) | #5536 | Cell Signaling Technology |
| **7.** | AMPKalpha | #5831 | Cell Signaling Technology |
| **8.** | Phospho-AMPKalpha (Thr172) | #2535 | Cell Signaling Technology |
| **9.** | HIF-1beta/ARNT | #5537 | Cell Signaling Technology |
| **10.** | HIF-1alpha | #36169 | Cell Signaling Technology |
| **11.** | HIF-2alpha | #59973 | Cell Signaling Technology |
| **12.** | Akt (pan) | #4691 | Cell Signaling Technology |
| **13.** | Phospho-Akt (Ser473) | #4060 | Cell Signaling Technology |
| **14.** | PKC alpha | #59754 | Cell Signaling Technology |
| **15.** | Phospho-PKC alpha/beta II (Thr638/641) | #9375 | Cell Signaling Technology |
| **16.** | CK2 alpha | #2656 | Cell Signaling Technology |
| **17.** | P70 S6 Kinase (S6K) | sc-8418 | Santa Cruz Biotechnology (Dallas, Texas, USA) |
| **18.** | Phospho-p70 S6 Kinase (pS6K) (Thr389) | #9234 | Cell Signaling Technology |
| **19.** | S6 Ribosomal Protein | #2217 | Cell Signaling Technology |
| **20.** | Phospho S6 Ribosomal Protein (Ser235/236) | #4858 | Cell Signaling Technology |
| **21.** | CA9 | #5649 | Cell Signaling Technology |
| **22.** | Ubiquitin (P4D1) (pan) | sc-8017 | Santa Cruz Biotechnology |
| **23.** | HA-tag | sc-7392 | Santa Cruz Biotechnology |
| **24.** | PGAM1 | #12098 | Cell Signaling Technology |
| **25.** | Hexokinase 2 (HK2) | #2867 | Cell Signaling Technology |
| **26.** | PKM2 | #4053 | Cell Signaling Technology |
| **27.** | Aldolase-A (ALDOA) | #8060 | Cell Signaling Technology |
| **28.** | SPHK1 | #12071 | Cell Signaling Technology |
| **29.** | SPHK2 | #32346 | Cell Signaling Technology |
| **30.** | SPTLC1 | 15376-1-AP | ProteinTech (Rosemont, Illinois, USA) |
| **31.** | SPTLC2 | 51012-2-AP | ProteinTech |
| **32.** | Hsp90 | #4877 | Cell Signaling Technology |
| **33.** | Hsc70 | sc-7298 | Santa Cruz Biotechnology |
